# A common cortical basis for variations in visual crowding

**DOI:** 10.1101/2023.12.07.570607

**Authors:** John A. Greenwood, Katarina Jerotic, Joseph E. Danter, Rhiannon J. Finnie, D. Samuel Schwarzkopf

## Abstract

Peripheral vision is limited by crowding, the disruptive effect of clutter on object recognition. Crowding varies markedly around the periphery, with e.g. stronger performance decrements with increasing eccentricity and in the upper vs. lower visual field. Although a number of neural substrates have been proposed for crowding, none to date can explain the full pattern of these variations. Here we examine the effects of crowding on object *appearance*. These effects are central to many models of crowding, and also vary markedly, causing target objects to appear more similar to flanker objects (assimilation) in some instances and dissimilar (repulsion) in others. We took 3 manipulations known to vary crowded performance (flankers in the same vs. different hemifield, the upper-lower visual field anisotropy, and the radial-tangential flanker anisotropy) and examined whether the effects on appearance vary similarly. In all cases, manipulations that increased performance impairments also increased assimilative errors, e.g. flankers on the radial axis around fixation gave high threshold elevation and assimilation, with reduced elevation and repulsion errors for tangential flankers. These linked variations in performance and appearance are well described by a population-coding model of crowding that varies the weighted combination of target vs. flanker population responses. We further demonstrate that this pattern is inconsistent with crowding being driven by either the cortical distance between elements or receptive-field size variations on their own. Instead, using a series of models we show that crowding could be driven by receptive field overlap – the intermixing of the spatial distribution of target/flanker population responses. Crowding is strong (with high performance decrements and assimilative biases) when the degree of spatial overlap in population responses is high and reduced (with low threshold elevation reduced assimilation or repulsion) when these responses are separable.

## Introduction

The fundamental limit on our peripheral vision is *crowding*, the impairment to object recognition that arises in clutter (1, 2). The disruption from crowding varies markedly with the position of both the target and surrounding flanker elements across peripheral vision, e.g. increasing as the target becomes more peripheral (1, 3) and in the upper vs. lower visual field (4–6). Although a number of neural substrates have been proposed to explain crowding, most fall short in explaining the full pattern of these variations. To date, variations in crowding have predominantly been examined via object recognition *performance*. A central aspect of many models of crowding (7, 8) is the observation that crowded errors are not random but rather follow the *appearance* of flanker elements (9, 10). To better understand the source of these variations, we first examined whether manipulations that vary crowded performance deficits also vary changes in appearance. From these observations we construct a computational model of crowding to simulate its variations and in turn examine whether they could derive from a common cortical basis.

Crowding is typically observed as a decrement in recognition performance for a target surrounded by flanker objects, relative to the target in isolation (1, 3). The magnitude of this performance decrement varies according to the separation between target and flanker elements, as well as the location of the target within the visual field. The most prominent of these variations is the increase in the magnitude of crowding with eccentricity (1, 3). Performance impairments are also stronger in the upper visual field than the lower, in the left hemifield vs. the right, and along the vertical meridian of the visual field than along the horizontal meridian (4–6). The location of the flankers also matters – those positioned along the radial axis with respect to fixation are more disruptive than those along the iso-eccentric tangential dimension (11), while more eccentric flankers will also produce greater crowding than flankers closer to fixation (4). Flankers positioned in the same visual hemifield as the target have also been found to produce more crowding than those presented to either side of the vertical meridian (12).

Beyond these performance decrements, a central aspect of many crowding models is the observation that crowding changes the appearance of the target. For instance, errors in the judgement of a target letter are more likely to match the flanker letters than a random letter (9, 13, 14). Systematic errors have also been found for a range of featural judgements, where the appearance of the target in terms of its feature positions (15, 16), orientation (17), motion and colour (18) are all biased in the direction of surrounding flanker elements. These assimilation effects increase the similarity between a target and its surrounding context. Errors follow the average of the target-flanker features when elements are similar to one another (15), while larger target-flanker differences give errors that resemble a substitution of the flanker identities (17, 19). Both of these error types are well described by population-coding models where crowding occurs due to the unwanted combination of the population responses to each element (17, 18, 20, 21). The implication of these systematic errors is that we make errors in clutter not because we see nothing, but because the target appearance is changed. A key assumption is that the effects of crowding on performance and appearance are intrinsically linked, yet this remains untested.

The effect of crowding on appearance has similarly been found to vary around the visual field. In addition to the above assimilative effects of crowding, the presence of flankers can at times make targets appear more dissimilar to the flankers (22, 23). At eccentricities near to the fovea, errors of repulsion are more common, while at far eccentricities the very same target/flanker orientations can cause assimilation errors (24). This shift from repulsion near the fovea to assimilation as eccentricity increases appears to be a general pattern, with similar effects reported for both the tilt illusion (25, 26) and Ebbinghaus illusion (27). In the context of crowding, repulsive errors can be incorporated into population pooling models with inhibitory surrounds (18), but how the resulting variations in target appearance relate to the above variations in performance is at present unclear.

Could a common cortical factor drive these variations around the visual field, affecting both performance and appearance? Several candidate factors have been proposed, the most prominent of which is the *cortical distance* between target and flanker elements. Because retinotopic visual areas show cortical magnification, with an expanded representation of the visual field around the fovea relative to the periphery (28–30), target and flanker elements with a fixed distance in the visual field would shift closer together cortically as they moved into the periphery. The increase in crowded performance decrements with eccentricity has been attributed to this decrease in cortical distance (31–34). Variations in the effect of crowding on appearance have also been linked with cortical distance, with the predominance of repulsion at parafoveal eccentricities attributed to the larger cortical distance between elements compared to the periphery, where the decrease in cortical distance promotes assimilation (24).

Variations in crowding strength have alternatively been linked with the spatial properties of receptive fields in visual cortex. Most prominent is the proposal that the pooling of target and flanker elements occurs when both fall in the same receptive field (35), suggesting that it is variations in *receptive field size* that drive the variations in crowding strength. fMRI estimates of the population receptive field (pRF) sizes in area V2 have accordingly been found to correlate with individual differences in crowded performance impairments (36). A third proposal is that crowding may derive from the *receptive field overlap* between adjacent neurons and their responses to the target and flanker elements (21, 37). Here, crowding arises when neurons responding to the target overlap in their spatial selectivity with neurons responding to the flankers, leading to the pooling of their signals at the population level. These three factors – cortical distance, receptive field size, and receptive field overlap – are obviously inter-related, given for instance the correlation between cortical magnification and pRF size (38), and that an increase in receptive field size would also increase their overlap. Each factor nonetheless makes divergent predictions for the myriad variations in crowding (as we examine later), making their respective role(s) in driving these variations unclear.

Altogether, in peripheral vision the presence of flankers around a target produces both performance impairments and systematic changes in target appearance. These effects vary with the location of target and flanker elements in the visual field. It is however unclear whether any of the cortical factors proposed to underlie crowding – cortical distance, receptive field size, and receptive field overlap – can explain the entirety of these variations. Here we examine three manipulations of target-flanker location, each with distinct predictions for these candidate factors, and measure their effects on performance and appearance. We first vary the hemifield location of target and flanker elements, following prior observations that crowding is stronger for elements presented on the same vs. different sides of the vertical meridian (12). The second is the upper-lower anisotropy, where crowding is stronger in the upper than the lower visual field (4). The third is the radial-tangential anisotropy (39), where flankers along the radial axis with respect to fixation cause greater crowding than those on the tangential axis. These three manipulations have thus far only been examined with respect to performance variations – if crowding is driven by a common cortical factor, then the effects of crowding on appearance should vary similarly. To examine whether a common cortical factor could explain all three effects, we then developed a series of models to simulate these variations. We demonstrate that the effects of crowding on performance and appearance do indeed co-vary, and that these effects are well captured by a population pooling model, but that cortical distance and receptive field size on their own are insufficient to drive these variations. Rather, we suggest that receptive field overlap is the most likely property that could serve as the common cortical factor for variations in crowding.

## Results

### Experiment 1: Left/right hemifield effects

Given the prominence of cortical distance as an explanation of variations in crowding (31–34), we first sought to dissociate physical from cortical distance using the division of the left and right sides of the visual field, which project to the right and left hemispheres, respectively (30, 40). Liu, Jiang (12) proposed that this hemispheric separation would cause a contralateral flanker (on the opposite side of the vertical meridian to the target) to be anatomically segregated and thus more distant cortically than an ipsilateral flanker (on the same side of the meridian), despite both having matched physical distance. Consistent with a role for cortical distance, percent-correct performance was found to be more disrupted by the ipsilateral flanker than the more cortically distant contralateral flanker.

In Experiment 1, we adapted the approach of Liu, Jiang (12) to also examine crowded effects on appearance. A target Gabor was presented 15° in the upper visual field, 1° to the left or right of the vertical meridian. The target was either presented alone, or with a single flanker that was either on the opposite side of the meridian, again 1° from the midline, or in the same hemifield (Figure 1A). Both flankers had the same physical separation from the target (2°). Participants judged whether the orientation of the target grating was clockwise (CW) or counter-clockwise (CCW) of vertical. If crowding depends on the cortical distance between target and flanker elements, then in addition to the effects on performance noted by Liu, Jiang (12), we should also observe that the cortically close ipsilateral flankers produce more assimilation than the cortically distant contralateral flankers.

**Figure 1.**
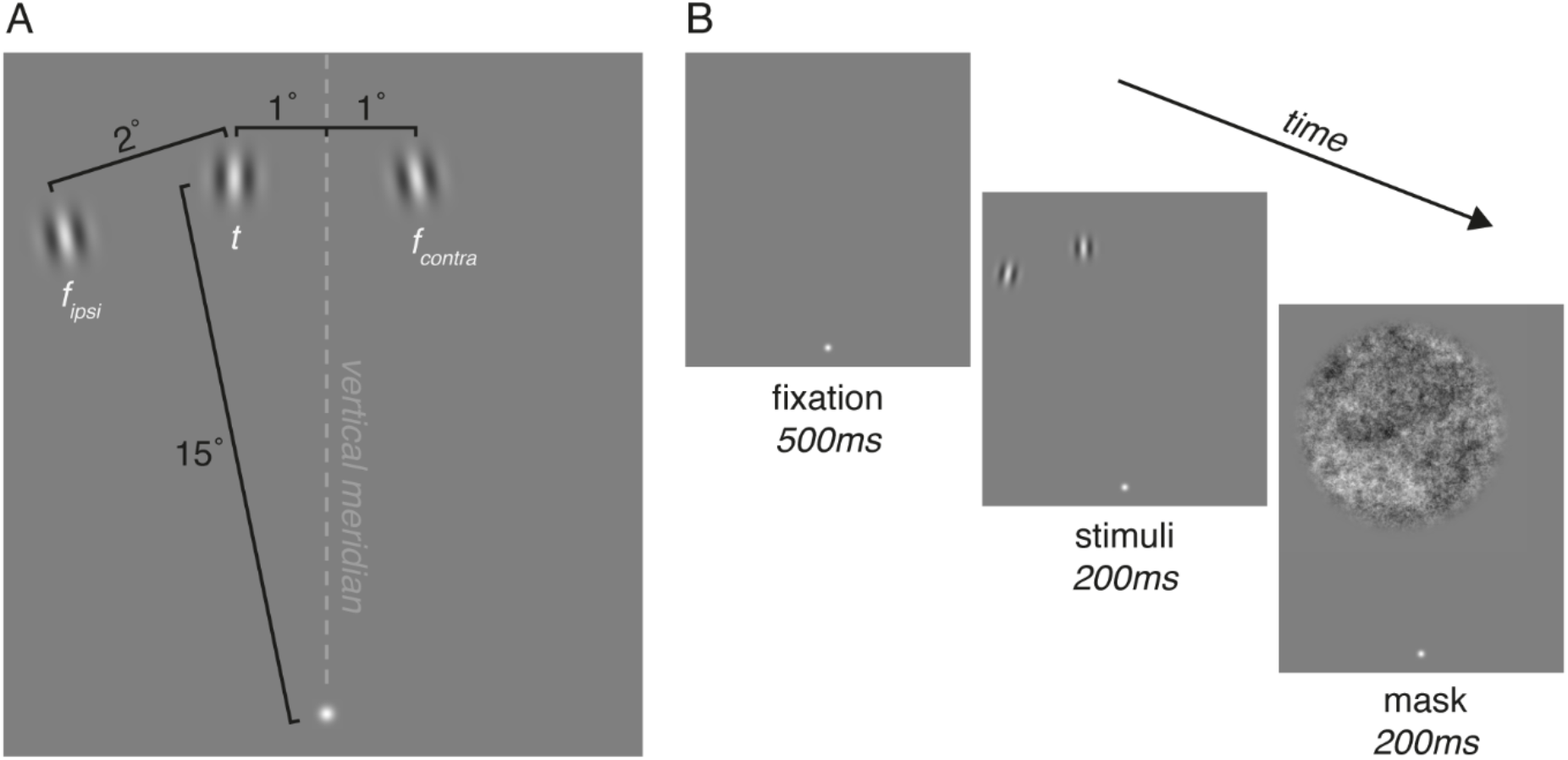
Stimuli and procedures for Experiment 1. **A.** Schematic of the stimulus arrangement. Participants fixated on a Gaussian blob, with a target Gabor (*t*) presented at 15° eccentricity, 1° from the vertical meridian (either to the left or right). The target was either presented in isolation, or with one flanker at a centre-to-centre distance of 2°, either in the same (*f_ipsi_*) or opposite hemifield (*f_contra_*). **B.** The time course of an example trial. Participants fixated for 500 ms prior to the presentation of stimuli for 200 ms. An example trial with a target in the left hemifield and an ipsilateral flanker is depicted. A 1/f noise mask followed for 200 ms.

Figure 2A shows example data from one participant in three conditions, each plotted as the proportion of CCW responses and fit with psychometric functions. Unflanked responses (grey) show a rapid transition between predominantly clockwise and counter-clockwise responses (indicating good performance) with a transition point close to vertical (indicating low bias). With a CCW-oriented flanker (red) in the contralateral visual field, the slope of the psychometric function is shallower (i.e. performance is worse), with an overall increase in CCW responses that shifts the midpoint of the function leftwards (indicating an assimilative bias). A similar slope is seen with the CW flanker (yellow), which decreases the rate of CW responses, shifting the midpoint rightwards (again indicative of assimilation). Altogether, the presence of a flanker both decreases performance and introduces biases consistent with a change in appearance.

**Figure 2.**
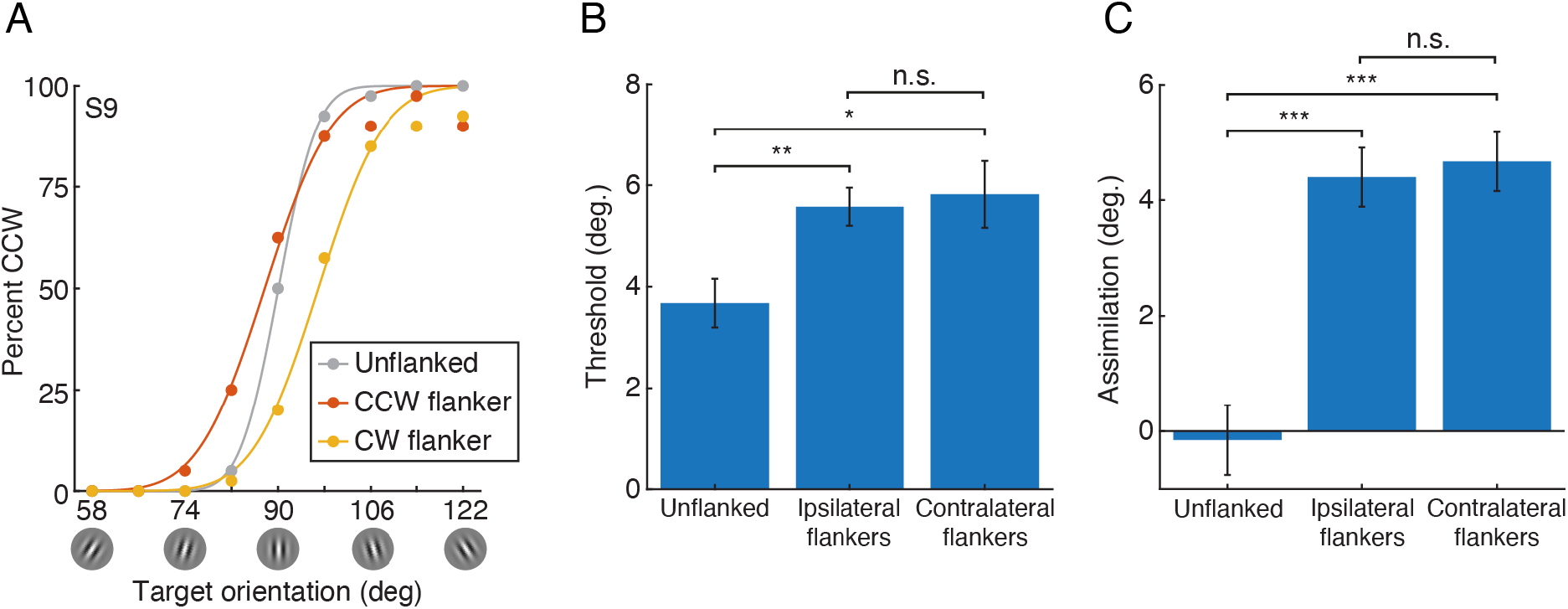
Data from Experiment 1. **A.** Example psychometric functions from one participant (S9). The proportion of CCW responses is plotted as a function of target orientation with an unflanked target (grey), and with flankers oriented −10° CW of vertical (yellow) and 10° CCW (red). Lines show the best-fitting psychometric function for each condition. Data is taken from conditions with the target in the right visual field, and for flanked conditions with a contralateral flanker (i.e. in the left visual field). **B.** Mean thresholds across participants (n = 10), separately for the three flanker conditions: unflanked, or with one flanker in either the ipsilateral or contralateral hemifield. Error bars denote ±1 standard error of the mean (SEM) across participants. Brackets indicate significance, with *** = p<.001, ** = p<.01, * = p<.05, n.s. = not significant. **C.** Mean assimilation values across participants, where positive values indicate assimilation and negative values repulsion. Plotted as in panel B.

To quantify performance, thresholds were taken as the difference in orientation required to shift performance from the midpoint to 75% CCW responses. Biases were measured as the orientation value at which the psychometric function reached its midpoint at 50% CCW responses. Values were subtracted from the reference orientation, with the sign of counter-clockwise biases reversed so that positive values indicated assimilative errors and negative values repulsion. Both thresholds and assimilation scores were then averaged across both target location conditions (left/right of the vertical meridian) and flanker orientation (CW/CCW of the target), both of which were found to have no effect in preliminary analyses. This left values for unflanked performance and with an ipsilateral vs. contralateral flanker for each participant.

Mean thresholds are shown in Figure 2B. Thresholds rose from around 4° when unflanked to 6° in the presence of a flanker. Paired-samples t-tests show that this crowding effect was significant for both ipsilateral (*t*_9_ = −4.73, *p* = .001) and contralateral (*t*_9_ = −2.68, *p* = .025) flanker conditions. However, the difference between ipsilateral and contralateral conditions was not significant (*t*_9_ = −0.51, *p* = .63). The presence of a single flanker thus induced a small but significant crowding effect, with both flanker locations causing an equivalent degree of crowding. The latter null result was not driven by outliers – on an individual level, 5/10 participants showed greater crowding in the ipsilateral condition, with the remainder showing the opposite.

Assimilation scores varied similarly, as shown in Figure 2C. On average, participants were unbiased with an unflanked target. The addition of a flanker increased responses that followed the flanker orientation, which required an average rotation around 4° to counteract. These assimilative errors were significantly different from unflanked biases for both the ipsilateral (*t*_9_ = −6.31, *p* < .0001) and contralateral (*t*_9_ = −6.15, *p* < .0001) conditions. Again however, the bias induced by the ipsilateral and contralateral flankers did not differ significantly (*t*_9_ = −0.61, *p* = .56). This effect was again not driven by outliers – if anything, individual values tended towards greater assimilation with the contralateral flanker (for 7/10 participants).

Our results do not replicate those of Liu, Jiang (12), where ipsilateral flankers were found to impair the percent-correct recognition of oriented Gabors more than contralateral flankers. We thus conducted power analyses to consider the reliability of our null result. If we consider the maximal crowding effect reported by Liu, Jiang (12) to be the difference between thresholds in the unflanked and ipsilateral flanker conditions, then the shift to contralateral flankers reduced this crowding effect by 57.6%. Applying this factor to our data, our sample (slightly larger than that of 12) had a probability of 77.88% to detect an effect of this size for the thresholds that we observe and a probability of 99.85% to detect an equivalent effect on assimilation. We also observe clear crowding effects in our data – both threshold elevation and assimilation increased with ipsilateral flankers, relative to unflanked performance. Both measures had large effect sizes (*d* = 1.48 and 2.81 for thresholds and assimilation, respectively), which even rose slightly with the contralateral flankers. We conclude that our null result is unlikely to derive from a power issue.

Two factors may explain this null effect. First, although our effects were clear, the observed effects of crowding were somewhat small, likely related to the presence of only a single flanker. Since an increase in flanker number can greatly increase crowding magnitude (3, 41), it is possible that the effect of cortical distance may become more evident for manipulations with a greater number of flankers. Second, although this hemifield manipulation may alter the cortical distance between the centroid locations of stimuli, it is likely less successful at altering the degree of receptive field overlap between the elements. That is, although the 1° distance from the vertical meridian may shift elements into distinct hemifields, the considerable size of receptive fields in peripheral vision could lead to a degree of overlap that nonetheless maintains the interference between these elements, particularly for neurons higher in the visual hierarchy (42, 43).

Importantly, despite this null result for cortical distance, there is a clear linkage between the effect of crowding on performance and appearance – the presence of a flanker element induced both an increase in thresholds and in assimilative bias. These results demonstrate that crowding is clearly occurring, with alterations to both performance and appearance, but that it is not being modulated by whether the flanker is located within the same or opposite hemifield. We next sought to increase the strength of these effects to better measure the nature of their co-variation.

### Experiment 2: The upper-lower field anisotropy

A second factor known to alter the strength of crowding is the upper-lower anisotropy. Several studies have observed larger performance decrements in the upper than the lower visual field (44, 45), along with larger interference zones in the upper field (4–6). This mirrors the more general performance anisotropies seen for tasks ranging from contrast sensitivity to illusory contour perception (46–51).

Several cortical factors could give rise to this difference in crowding between the upper and lower fields. Cortical distance could play a role, given evidence from both physiological measurements (52, 53) and neuroimaging (54, 55) that the surface area of V1 and V2 is smaller in the upper field than the lower. The smaller surface area in the upper field would cause flankers at a given physical distance to be closer on the cortical surface than elements with the same target-flanker distance presented in the lower visual field (45). Receptive fields have also been found to be larger and more elliptical in the upper than the lower field (56), which could similarly account for the differences in crowding. These changes in receptive field size would also alter receptive field overlap, as has been argued for the inner-outer anisotropy (21, 37).

The effect of the upper-lower anisotropy on appearance is unknown. As above, if crowded effects on performance and appearance are linked, then assimilation should be stronger in the upper field than the lower. We tested this possibility in Experiment 2. Stimuli and procedures were similar to those of Experiment 1, though with elements placed along the vertical meridian, either in the upper or lower visual field. The target was presented either in isolation or between two flankers along the radial dimension at 4 eccentricities in each field.

As before, psychometric functions were fit to individual conditions, with threshold and midpoint values taken as measures of performance and appearance, respectively. For unflanked conditions, thresholds ranged from 1.8-2.2° on average, with no clear variation by either eccentricity or visual field. Average midpoints were within 0.5° of 90°, again with no variation by eccentricity or visual field. A two-way ANOVA run on these values confirmed this, with non-significant main effects for both eccentricity and visual field, and a non-significant interaction (all *F*<2). We attribute this to the high visibility of these elements given the approximate M-scaling of stimulus size and spatial frequency as a function of eccentricity.

Flanked conditions were again pooled across the flanker orientation conditions (clockwise/counter-clockwise). Threshold elevation scores were calculated for each participant by dividing crowded thresholds by the mean unflanked threshold (pooled across eccentricities and visual fields given the lack of variation). Mean threshold elevation values are plotted in Figure 3A. While the manipulations of Experiment 1 produced elevations around 1.5 times unflanked thresholds, here threshold elevation peaks around 10 times unflanked performance. Threshold elevation increases with eccentricity, and is greater in the upper visual field than in the lower. These values were submitted to a three-way mixed effects ANOVA, with eccentricity and visual field as fixed effects, and participant as a random effect. Significant main effects were obtained for eccentricity *F*(3, 21) = 13.94, *p* < .001 and visual field *F*(1, 21) = 13.80, *p* = .008, but not for participants, *F*(7,21) = 2.93, *p* = 0.09. The interaction between eccentricity and visual field was also significant, *F*(3,21) = 4.80, *p* = .011, given the steeper rise in threshold elevation with eccentricity for the upper than the lower field. The interaction between eccentricity and participant was not significant (F<1), though the interaction between visual field and participant was significant, *F*(7,21) = 6.27, *p* = .0005.

**Figure 3.**
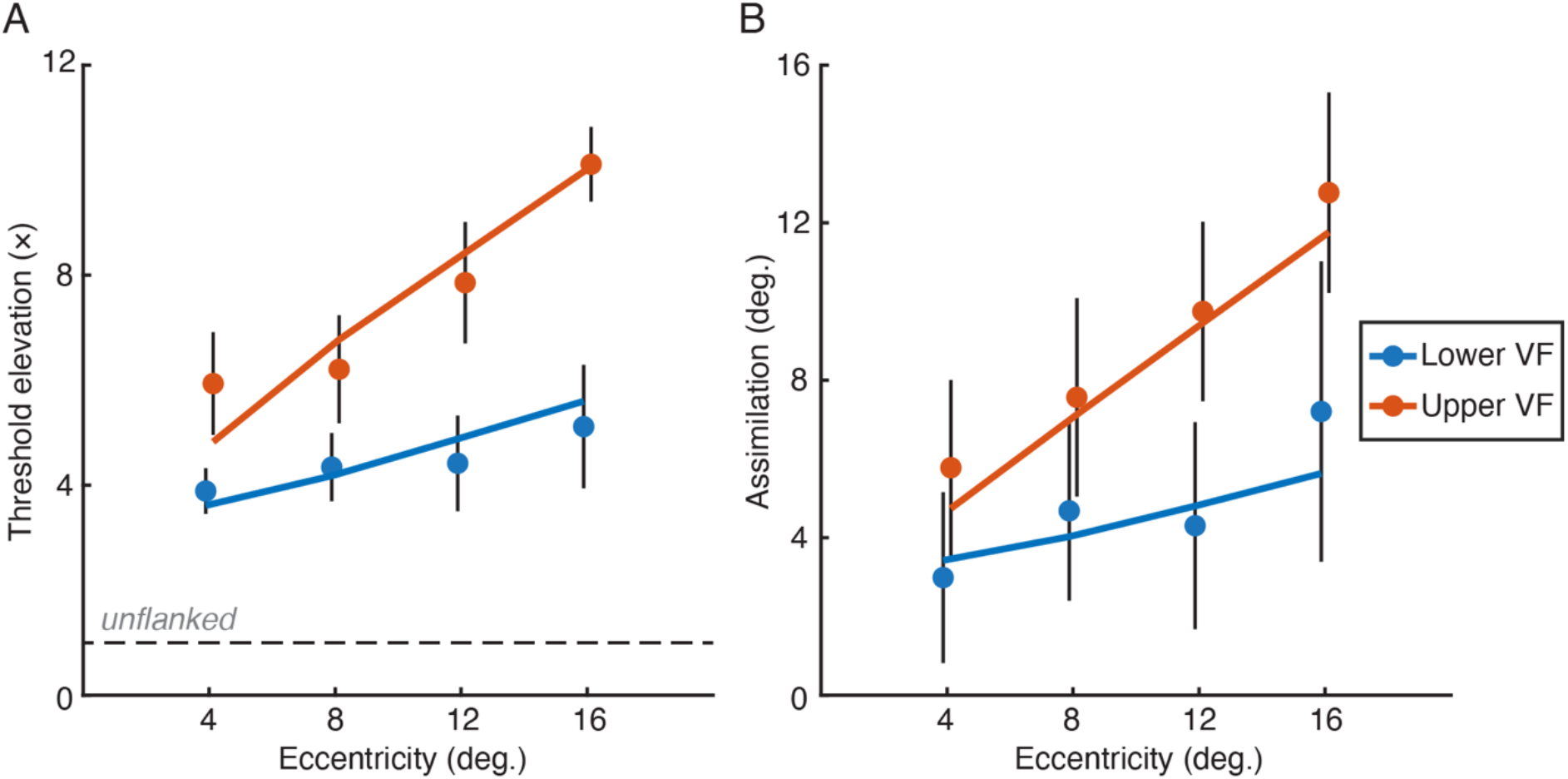
Results from Experiment 2. **A.** Mean threshold elevation scores (flanked thresholds divided by unflanked) as a function of eccentricity, separately for the lower (blue) and upper (orange) visual fields. Unflanked performance is indicated via the dashed black line. Points show the mean across participant, with error bars showing the SEM. Lines show the output of the best-fitting population pooling model. Data have been offset slightly on the x-axis for clarity. **B.** Mean assimilation scores plotted as a function of eccentricity, where positive values indicate assimilation and negative values repulsion, plotted as in panel A.

A similar pattern is evident for assimilation scores (Figure 3B). Our target-flanker configuration produced predominantly assimilative biases, which rose with eccentricity and were greater overall in the upper than the lower field. A three-way mixed effects ANOVA was again conducted, with significant main effects for eccentricity *F*(3, 21) = 7.61, *p* = .001, visual field *F*(1, 21) = 6.47, *p* = .038, and participant *F*(7,21) = 6.85, *p* = 0.005. Here the interaction between eccentricity and visual field was not significant, *F*(3,21) = 2.21, *p* = .117, though the interactions between eccentricity and participant, *F*(21,21) = 2.69, *p* = .014, and between visual field and participant *F*(7,21) = 10.02, *p* < .001, were both significant.

We demonstrate here that the upper visual field shows both stronger performance impairments from crowding and a higher level of assimilative biases relative to the lower visual field, even for elements at matched eccentricities. Our threshold elevation values replicate prior demonstrations of greater performance decrements in the upper field (44, 45). The rise in assimilation with eccentricity also replicates the findings of Mareschal, Morgan (24), though this is the first observation that these effects are magnified in the upper field. As above, these effects could be driven by the cortical separation between elements, changes in receptive field size and/or receptive field overlap.

The strength of the assimilation effects observed in our results was somewhat unexpected, given the repulsive biases observed at closer eccentricities by Mareschal, Morgan (24). This may be due to the closer target-flanker separations in the present study (e.g. 1.2° at 4° eccentricity in our study vs. 1.8° in Mareschal *et al*). Given their observation that target-flanker separation can alter the balance between assimilation and repulsion, repulsive effects might emerge with larger target-flanker separations. In Experiment 3 we thus sought to separate the effects of cortical distance vs. receptive field size and overlap, whilst also including variations in target-flanker separation to induce a wider range of both repulsive and assimilative biases.

### Experiment 3: The radial-tangential anisotropy

One of the most consistently observed variations in the strength of crowding is the radial-tangential anisotropy – flankers along the radial axis with respect to fixation produce stronger performance decrements and larger interference zones than those along the tangential axis (4, 5, 39, 57). The effect of this anisotropy on appearance is unknown. If the effects of crowding on appearance follow those for performance (as they have in Experiments 1 and 2), then we predict greater assimilation for radial than tangential flankers.

The radial-tangential anisotropy has been attributed to several cortical factors. As with the two preceding variations, cortical distance could produce these effects (34). However, the representation of the visual field along the radial axis extending outwards from the fovea shows less compression (due to cortical magnification) than the representation along the tangential or polar angle axis (54, 58). Were cortical distance alone to determine the strength of crowding, this would predict greater effects on the tangential axis – the opposite of the radial-tangential anisotropy. Alternatively, receptive field shape may play a role: an elongation of receptive fields along the radial axis would increase the likelihood of flankers falling within the same receptive field relative to those on the tangential axis. This elliptical shape has indeed been observed in macaque V4 (35), potentially driven by a constant sampling of V1 inputs in conjunction with the effect of cortical magnification. Similar levels of radial elongation have been observed for pRF measurements in human cortex (56), though this is not found in other analyses (59). Finally, these anisotropies in receptive field shape would also modulate receptive field overlap (21, 37) in a direction consistent with the anisotropy.

Stimuli and procedures were similar to those of Experiment 2, with target orientation judgements measured in the upper visual field, either for an isolated target or with two flankers on either the radial or tangential axis. Flankers were placed at one of several target-flanker separations (from 1.6-3.2°). Psychometric functions were again fit to each condition for each participant. When unflanked, thresholds were on average 2.15°, with a midpoint of 90.1°, similar to Experiment 2. Threshold elevation values were then calculated for the flanked conditions, with means plotted in Figure 4A, separately for each target-flanker separation and axis. Close target-flanker separations produced the most threshold elevation, which decreased as the flankers were spaced apart. This was particularly so for the radial axis where threshold elevation was approximately twice as high as tangential flankers at each separation. This pattern is borne out by a three-way mixed-effects ANOVA, with significant main effects for axis, *F*(1,28) = 20.13, *p* = .003, and separation, *F*(4,28) = 29.50, *p* < .001. The interaction between separation and axis was not significant, *F*(4,28) = 1.97, *p* = 0.12. The main effect of participants was not significant, *F*(7,28) = 1.92, *p* = .21, nor was the interaction between separation and participants, F<1, though the interaction between axis and participant was significant, *F*(7,28) = 14.26, *p* <.001.

**Figure 4.**
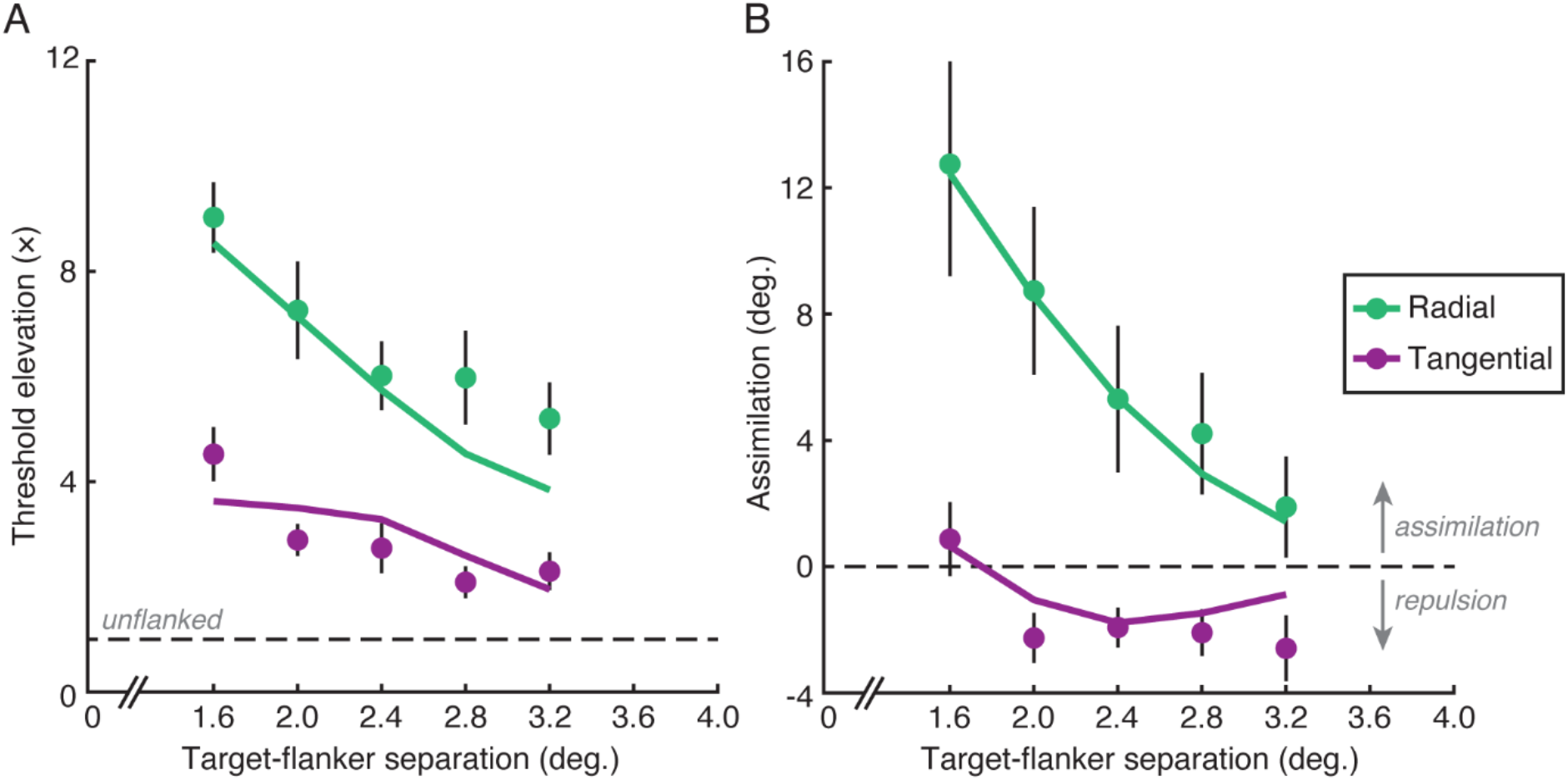
Results from Experiment 3. **A.** Mean threshold elevation scores as a function of target-flanker separation, separately for flankers on the radial (green) and tangential (purple) axes. Unflanked performance is shown via the dashed black line. Points show the mean across participants, with error bars showing the SEM. Lines show the output of the best-fitting population pooling model. **B.** Mean assimilation scores plotted as a function of target-flanker separation, where positive values indicate assimilation and negative values repulsion (separated by the dashed line). Plotting conventions as in panel A.

Assimilation scores were also calculated, as before (Figure 4B). Here too assimilation was highest at closer target-flanker separations, and greater with radial flankers than tangential. Interestingly, biases with tangential flankers show assimilation at the closest separation, which transitions to repulsion at larger separations. This pattern is again supported by a three-way ANOVA, with significant main effects for axis, *F*(1,28) = 9.86, *p* = .016, and separation, *F*(4,28) = 15.84, *p* < .001. The main effect of participants was not significant, *F*(7,28) = 1.08, *p* = .47. The interaction between separation and axis was significant here, *F*(4,28) = 3.29, *p* = 0.025, as was the interaction between axis and participant, *F*(7,28) = 11.23, *p* <.001. The interaction between separation and participants was not significant, F<1.

One issue is that the radial-tangential anisotropy observed here is confounded with the alignment of our elements. Because the Gabor orientations were centred on vertical, a change from radial to tangential flankers also shifted orientations from being collinear to parallel, which is known to modulate crowding (23, 60, 61). We thus ran an additional experiment, with elements rotated by 90° to reverse the nature of the collinearity (described in Appendix A and Figure S1). This manipulation gave the same pattern of results, confirming that it is the spatial location of the flankers that drives this effect rather than their respective orientations.

Our observation that radial flankers produce twice the threshold elevation of tangential flankers replicates the well-known radial-tangential anisotropy (39). Here we show that this anisotropy also affects appearance, with radial flankers producing greater assimilation than tangential flankers. The latter also began with assimilative biases at the closest separations, shifting to repulsion as target-flanker separation increased. As outlined above, the increased threshold elevation and assimilation with radial (vs. tangential) flankers is inconsistent with the most straightforward relationship between cortical distance and assimilation. Factors including receptive field size and overlap would thus appear to be better explanations for these variations, which we next examined with a series of models.

## Modelling

All three experiments show co-variation between threshold elevation and assimilation. These values are indeed correlated (Figure S2 of Appendix B). Given this co-variation, and the modulation of these values by both the upper-lower and radial-tangential anisotropies (Experiments 2 and 3), we next sought to model these factors and explore the potential for a common cortical basis. We began by modelling the effects of crowding on appearance and performance, which we then used as a basis to examine the cortical factors that may drive these variations.

### Population pooling model of crowding

As outlined in the introduction, the systematicity of crowded errors is well explained by ‘pooling’ models of crowding (7, 8, 15, 17), which model these effects as a combination of the target and flanker elements. Population pooling models are particularly well equipped to account for the diverse effects of crowding on appearance, given prior simulations of both assimilation and repulsion errors (17, 18, 21), as well as crowding effects in typical vision and amblyopia (20). Here we examine whether population pooling can account for the variations in crowding found for both threshold elevation and assimilation in the current study. Given the lack of modulation of these factors in Experiment 1, we focus on Experiments 2 and 3.

Full details of the model are outlined in the *Materials & Methods*. In brief, the model simulates the response of a population of detectors, each with a Gaussian sensitivity profile for orientation and an inhibitory surround. Population responses were determined for target and flanker orientations separately and combined via weights, as in recent models (18, 20). Variations in crowding were produced by varying these weights from 0-1. The peak of the resultant population responses gives an estimate of perceived orientation on each trial, which was used across conditions to generate proportion correct responses, with psychometric functions fit to give midpoint and threshold values.

For Experiment 2, the output of the best-fitting population pooling model is plotted against the data in Figure 3. The model captures the rise in threshold elevation with eccentricity (Figure 3A) through an increase in flanker weights, which increases both the level of noise introduced into the population by the flanker population response and the number of trials with a peak response pulled in the direction of flankers. A steeper slope for the upper field captures the increased threshold elevation relative to the lower field. For biases (Figure 3B), these variations in weights similarly capture both the rise in assimilation with eccentricity and the increased assimilation in the upper field. A population pooling process can thus produce variations in both threshold elevation and assimilation across the upper and lower fields.

To simulate the results of Experiment 3, flanker weights were varied as a function of target-flanker separation. For threshold elevation (Figure 4A), the model captures the drop in threshold elevation with increasing target-flanker separation, as well as the greater degree of threshold elevation along the radial vs tangential axis. Figure 4B shows the biases, where the model similarly is able to capture the high degree of assimilation for radial flankers and their decline with increasing separation, as well as the transition from assimilation to repulsion on the tangential axis. As with the upper-lower anisotropy, a population pooling process can therefore simulate the effects of crowding on threshold elevation and biases, here including errors of both assimilation and repulsion.

### Modelling the common cortical factor

The population pooling model demonstrates that variations in crowding can be simulated by varying the weights applied to the pooling of target and flanker elements in a common population. We can therefore reduce the effect of crowding on two factors (performance and appearance) down to one: crowding varies because the weighting applied to the flanker response varies. This in turn raises the question: what drives the variations in flanker weights?

At this point, we can exclude cortical distance as a likely common factor for these variations. In Experiment 1, we did not observe variations in crowding for elements presented to the same vs. different hemifields, as others have predicted for the effect of cortical distance (12). Similarly, in Experiment 3, cortical distance variations make the opposite prediction (54, 58) to the radial-tangential anisotropy for both appearance and performance. Receptive field size may be a more likely contender, particularly for the upper-lower anisotropy, given observations that pRFs are larger in the upper than the lower visual field (56). However, for receptive field size/shape to account for the radial/tangential anisotropy, elliptical receptive field shapes would be required (56), which has been contested (59), suggesting either that fields are circular or that the aspect ratio is small (and difficult to measure).

We can however unify these observations using *receptive field overlap* – the degree to which the target and flanker activate common neurons via the overlap in their receptive fields. To better fit with the population pooling process described above, we can further consider receptive field overlap on a population level – the extent to which the population responses to the target and flanker elements activate the same neurons/detectors. We thus sought to quantify receptive field overlap using the responses of a population of model receptive fields to stimuli presented in the visual field. In this model, each detector is a two-dimensional Gaussian element, with the location of our stimuli similarly represented by Gaussian elements, given the shape of their contrast envelopes. The response of each detector to the target and flanker stimuli was obtained by convolving its receptive field profile with that of the stimulus and taking the sum (separately for the target and flankers). In this way, responses were determined by the spatial overlap between each detector’s receptive field and the stimulus. To determine the spatial distribution of the population response to our stimuli, each detector contributed its own receptive-field profile to the final population response to the stimulus – a detector with a small receptive field would produce a tightly distributed estimate of the target location, while a larger receptive field would produce a broader estimate. The magnitude of this Gaussian spatial profile was given by multiplying the Gaussian receptive field by the magnitude of the overlap between the receptive field and the stimulus. We can then quantify the degree of this overlap by multiplying the two images together and taking the sum.

To illustrate the operation of the model, we begin with a simple example with a 3×3 grid of neurons in the upper visual field in order to examine the variations necessary to produce a radial-tangential anisotropy. Note that receptive field overlap is sensitive to several factors including receptive field size and shape, the distance between the receptive fields in the visual field (likely related to their cortical distance), and the sampling density of neurons responding to a given part of the visual field (similarly related to cortical magnification). Consider a case where these receptive fields have centres that are equally spaced horizontally and vertically, with matched receptive field sizes (Figure 5A). With a target centred at 8° on the middle receptive field, and two flankers positioned either radially (above/below) or tangentially (left/right), these detectors give a population response to the target that is shown in green in Figure 5B, with the population response to flankers in purple. Regions where these signals overlap are shown in white. By multiplying these distributions together and taking the sum, we can quantify this overlap. Normalised values of this overlap sum (divided by the response to the radial flankers) are presented in Figure 5C. Because the receptive fields are of constant size and separation in this example, there is no difference between radial and tangential axes.

**Figure 5.**
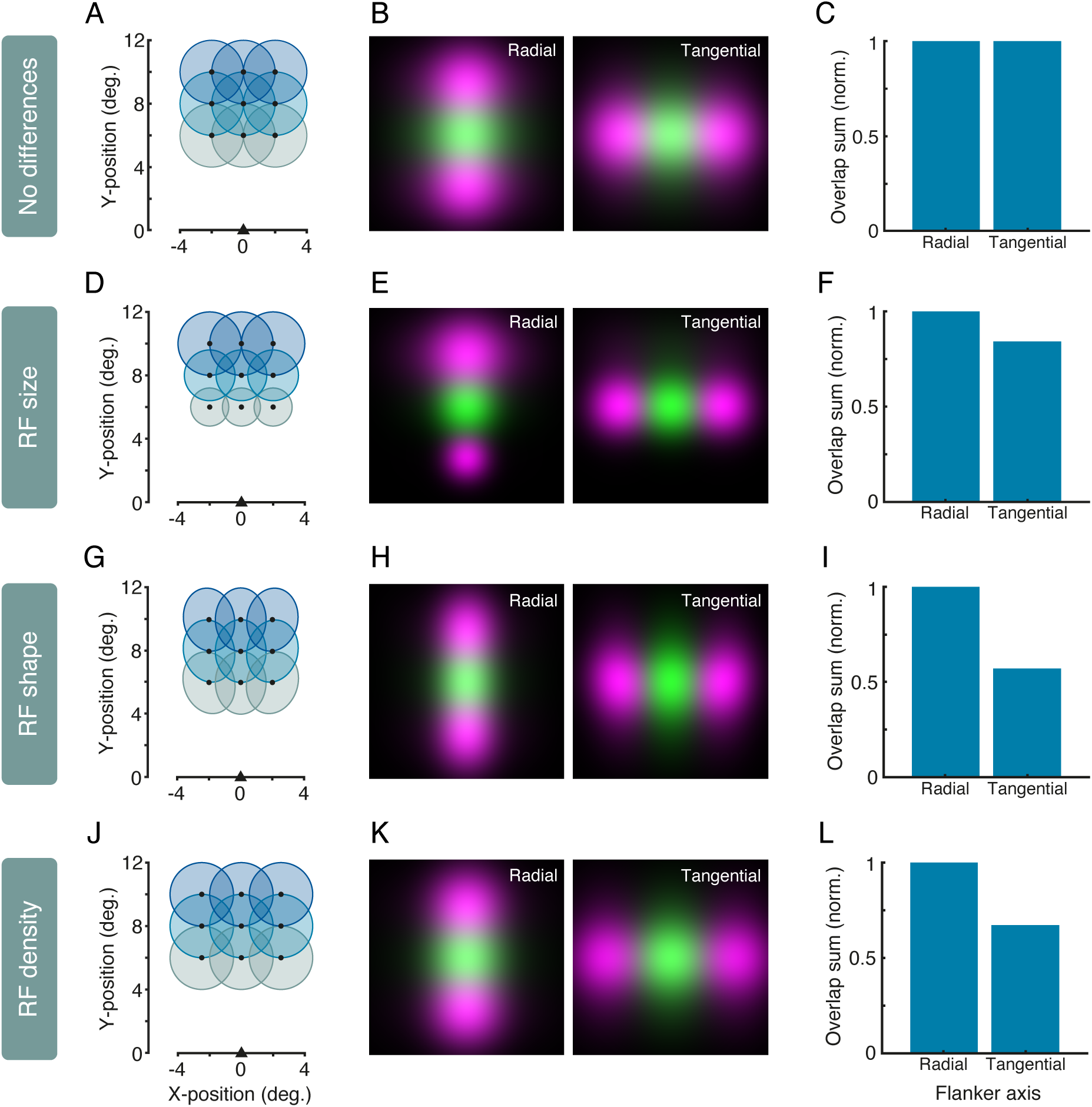
Receptive field overlap in a simple population of 9 detectors. The first row shows a population with circular Gaussian receptive fields of matched size and constant density/spacing in the horizontal and vertical dimensions (shown as circles in panel A). Panel B shows the response of these detectors to a target at 8 degrees in green, and two radially-positioned flankers (left) or tangential flankers (right) in purple, with regions of overlap shown as white. The sum of the overlap between these target and flanker responses is shown in panel C, with responses normalised by dividing by the radial overlap sum. The second row shows the same values for a population of detectors where receptive field sizes increase with eccentricity (panels D-F). In the third row, the detectors are elliptical, here with an aspect ratio of 1.25, oriented towards the fovea (panels G-I). The final row shows a population of circular Gaussians where the density/spacing on the horizontal dimension is 1.25 times larger than on the vertical dimension (panels J-L).

We can however alter several properties in this population that cause differences on the radial vs. tangential dimensions. The detectors in Figure 5D have receptive fields that increase in size with eccentricity (as each row progresses away from the ‘fovea’). In this case, the larger outwards receptive fields cause the spatial distribution of the radial flankers (Figure 5E, left panel) to overlap more greatly with the target than the distribution of the tangential flankers (right panel). This creates an anisotropy in the overlap sum (Figure 5F), with more overlap on the radial axis. A similar anisotropy can be obtained by altering the aspect ratio of the Gaussian receptive fields, such that each becomes an ellipse oriented inwards towards the fovea (Figure 5G). Here, with an aspect ratio of 1.25, the overlap sum is greater between the target response and that of the radial flankers (Figure 5H, left) than the tangential flankers (right), again creating an anisotropy in the overlap sum (Figure 5I). Finally, altering receptive field density by shifting the position of receptive fields (holding the above factors constant) can also alter the overlap sum. The population in Figure 5J have circular receptive fields with the same vertical positions as above, but with a separation on the horizontal axis that is 1.25 times that of the radial axis. This gives greater overlap between the target and the radial flankers (Figure 5K, left) than the tangential flankers (right), again causing an anisotropy in the overlap sum (Figure 5L). These simple demonstrations show that at least three factors can vary the degree of receptive field overlap – the size, aspect ratio, and density of receptive fields – which in turn could drive the strength of crowding.

We next used a larger population of detectors to examine whether receptive field overlap could simulate the upper-lower and radial-tangential variations in crowding observed above. Given that our simulations of crowded performance and appearance could be captured by varying the flanker weights in a population pooling model, here we ask whether receptive field overlap could in turn drive these weights. To do so, we fit the parameters of model populations of neurons using the sum of the receptive field overlap values to the flanker weights from the fits of the population pooling model in Experiments 2 and 3 (see *Materials & Methods* for details).

Given proposals that receptive field size is the key factor driving crowding (35, 36), we first examined whether this factor alone could alter receptive field overlap in a way that gives both the upper-lower and radial-tangential anisotropies. We began with the straightforward assumption that receptive field size increases with eccentricity, a property observed throughout visual cortex (42, 43) and which could produce the radial-tangential anisotropy in some circumstances (as in Figure 5D-F). Receptive field size was thus varied linearly with eccentricity. To simulate the upper-lower anisotropy of Experiment 2, we added a factor to multiply the slope of this function in the lower visual field, causing it to rise less steeply with eccentricity than in the upper field, similar to estimates from pRF mapping (51, 62).

The resulting population varied only in receptive field size, both with eccentricity and in the upper vs. lower visual field, as shown schematically in Figure 6A. These flanker weights from Experiment 2 are shown in Figure 6B, along with the best-fitting overlap sum values from the receptive field model. The pattern of receptive field overlap values clearly follows the pattern of the flanker weights. Namely, receptive field overlap increases with eccentricity, driven by the increase in receptive field sizes, with greater overlap in the upper than the lower visual field, driven by a more rapid rise in receptive field size in the upper visual field. Equivalent values for the radial-tangential anisotropy in Experiment 3 are shown in Figure 6C (NB. fit only to the positive weights in the population pooling model, see *Materials & Methods*). Here, although the population gave receptive-field overlap values that decreased as the spacing between target and flanker elements increased, overlap values did not differ on the radial and tangential axes in the same way as the flanker weights of the population pooling model. That is, although differences in receptive field size with eccentricity *can* produce a radial-tangential anisotropy, as shown in Figure 5, the parameters fit to the data of Experiments 2 and 3 gave an increase in receptive field size with eccentricity that was insufficient to produce the radial-tangential differences of Experiment 3. On their own, variations in receptive field size are thus unable to account for all of the variations in crowding.

**Figure 6.**
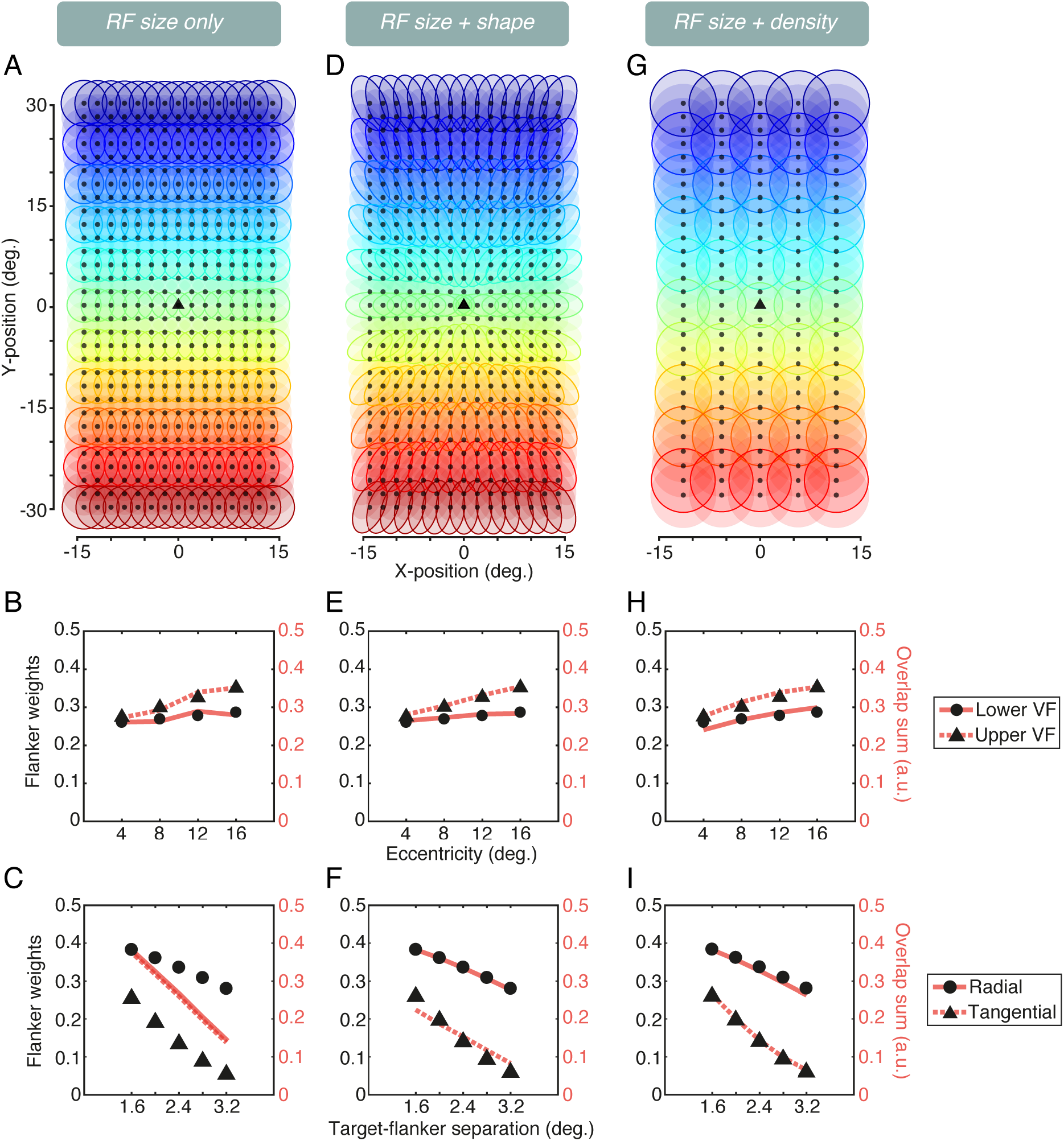
Simulations of receptive field overlap with three population models. The first model has receptive fields that tile the visual field, which vary only in their receptive field size (“RF size only”). **A.** A schematic representation of the receptive fields as they tile the visual field. Fixation is shown as a black triangle, with the centre of each receptive field shown as black dots. Receptive fields in each y-position are differently coloured for clarity, with selected rows outlined to show receptive field size (the full-width at half-maximum of each Gaussian element). **B.** Correspondence between the flanker weights for the population pooling model fit to the data of Experiment 2 (black points) and the receptive field overlap values fit to these weights (red lines). Values are plotted as a function of eccentricity, separately for the upper and lower fields. **C.** Correspondence between the flanker weights for the population pooling model of Experiment 3 (black points) and the best-fitting receptive field overlap values (red lines), plotted as a function of target-flanker separation, separately for the radial and tangential dimensions. **D-F.** Model details and output where the receptive fields also vary in their shape – becoming elliptical and oriented inwards towards the fovea. **G-I.** Model details and output where the receptive fields vary in their size (with eccentricity) and in their density/spacing between the upper-lower fields and on the radial vs tangential dimensions.

Following proposals that receptive field shapes may be elliptical rather than circular (56), we next adapted the aspect ratio of the receptive fields in the above model, orienting them such that their major axis aligned with fixation (depicted in Figure 6D). When fit to both experiments, the output of the model continued to give a good characterisation of the upper-lower anisotropy in Experiment 2 (Figure 6E), again driven by a more rapid rise in receptive field size with eccentricity in the upper field. For Experiment 3 (Figure 6F), the model fit improved substantially over that with receptive field size variations alone – now, in addition to the decline in overlap with increasing target-flanker separation, there is also a greater degree of overlap along the radial dimension than the tangential, driven by the elliptical shape of the receptive fields. The best-fitting model required an aspect ratio of 1.89 (radial:tangential) to produce this effect.

Given that evidence for elliptical receptive fields is mixed (59), we next examined whether variations in receptive field density could also give rise to the observed patterns. Here we fit a model with circular receptive fields that increased in size with eccentricity equivalently in all directions, without size variations between the upper and lower fields. Density was varied by changing RF separation, both in upper vs. lower fields and on the vertical (radial) vs. horizontal (tangential) dimensions. The best-fitting population is shown schematically in Figure 6G. Figure 6H plots the receptive field sum values of the best-fitting model against the model weights for Experiment 2 on the upper-lower anisotropy. A small difference in RF density between upper and lower fields was sufficient to produce this difference – the centre-to-centre distance between receptive fields was 1.08 times larger in the lower field than the upper field (i.e. detectors were more closely spaced in the upper field). The radial-tangential anisotropy could also be simulated by this population (Figure 6I). Here the horizontal (tangential) spacing of detectors was 2.83 times greater than that of the vertical (radial) spacing. A population of receptive fields that varies in their density in the upper vs. lower field and on the radial vs. tangential dimension (along with increasing RF size with eccentricity) can therefore produce a pattern of receptive field overlap that matches the variations in flanker weights in our population model. Ultimately, these models demonstrate that the overlap in the population response to target and flanker elements could be the common factor that drives crowding, itself driven by variation in factors including the size, shape, and density of receptive fields.

## General discussion

Crowding not only makes objects more difficult to identify but also alters their appearance. Here we show these effects are linked: both threshold elevation and assimilative effects on target appearance are greater in the upper visual field than the lower, and with flankers on the radial axis vs. the tangential axis (where appearance effects can also flip to become errors of repulsion). A population pooling model of crowding can account for all these effects through variations in the weights applied to the pooling process. This pattern of variations is inconsistent with them being driven solely by either cortical distance or receptive field size. Instead, we demonstrate with a model population of neurons that the degree of *receptive field overlap* could serve this function, with crowding increasing as the spatial response to the flankers overlaps with the response to the target element (driven by the size, shape, and density of RFs in the population). Greater receptive field overlap would cause the flankers to contribute more to the combined response than the target, increasing threshold elevation (through added noise) and the assimilative change in target appearance, matching the variations in flanker weights used in our population pooling model. Just as the effects of crowding are best captured by population-based processes, so too must we consider the population response to each element in considering the neural basis of these effects.

The observed coupling between performance and appearance in crowding is consistent with arguments for a general link between bias and discriminability in visual perception (63, 64). This coupling is also a central aspect of “pooling models” that depict crowding as the unwanted combination of target and flanker elements (7, 8, 15, 17). The generality of these models is evident in their simulation of the effect of crowding on motion and colour (18), higher-level elements such as faces (65), and associated elevations in clinical cases like amblyopia (20). A similar coupling between performance and appearance could also arise in the broader class of pooling-based “texturisation” models that depict crowding as the extraction of summary statistics across wide regions of the visual field (7, 66, 67). In contrast, models that do not account for this coupling cannot be said to provide a full account of crowding, most notably in the case of attentional (44, 68) and grouping accounts (69) that focus only on the disruption to performance.

Our instantiation of receptive field overlap as a common factor driving these variations differs from prior proposals (37) in that it incorporates the response of a large number of receptive fields that tile the visual field. As above, what we propose is more akin to a population receptive field overlap, in the same way that pooling models of crowding have been elaborated to incorporate populations of orientation detectors (17). We suggest that variations in receptive field overlap could provide the basis for the weighting values used in these population pooling models, with greater overlap between target and flanker response distributions leading to a stronger contribution of the flanker signals in the pooled response distributions. The unwanted combination of target and flanker population responses is also similar to physiological analyses linking crowding to integrative processes in area V4 (70), though neural correlates of crowding have been observed at several points in the human visual hierarchy (71).

Our modelling of receptive field overlap further suggests that there are multiple physiological factors that could alter this property to give the observed variations in crowding, including receptive field size, shape, and density. The utility of receptive field overlap is further evident in our observation that none of these factors *in isolation* can drive these variations. Receptive field size is an obvious contender in this regard, as proposed previously (35, 36). Given the well-established increase in the size of receptive fields with eccentricity, particularly in early visual areas (42, 43), this factor could drive the increase in crowding with eccentricity observed both here in Experiment 2 and many times previously (1, 3). Differences in receptive field size could also drive the upper-lower anisotropy, given observations that pRF sizes are larger in the upper field than the lower (51, 62). However, we show that the radial-tangential anisotropy cannot be produced by variations in receptive field size alone, at least within a population that can also reproduce the changes in crowding with eccentricity (Figure 6C).

Nonetheless, other factors could operate in conjunction with receptive field size to alter receptive field overlap. The first possibility is that receptive fields may be elliptical in shape, consistent with observations from pRF modelling (56) and single-cell recordings (35). The addition of this factor to variations in receptive field size allowed a pattern of receptive field overlap that matched both the radial-tangential and upper-lower anisotropies (Figure 6F). Although the reliability of elliptical pRFs has been disputed (59), it may be that small variations in this property (likely difficult to detect with neuroimaging) are sufficient to drive the observed differences, particularly if they operate in conjunction with other variations.

The overlap between target and flanker signals on the cortical surface could also be driven by the arrangement of receptive fields. Our models show that circular receptive fields that increase in size with eccentricity can reproduce all of the observed anisotropies when their density is higher in the upper vs. lower visual field and along the radial vs. tangential dimension with respect to fixation (Figure 6I). For the upper-lower difference this may relate to observations that the surface area of V1 is greater for the lower field than the upper (51, 72). The greater surface area of the lower field means that neurons would effectively be further away and likely more numerous, giving rise to the observed effects. The density and/or spacing of receptive fields in the tangential vs. radial dimension is harder to quantify.

One might expect a relationship between the density of RFs in the visual field and their separation on the cortical surface. We demonstrate here that cortical separation *on its own* is unable to account for the variations in crowding observed herein. Most clearly, in Experiment 1, we failed to replicate the results of Liu et al. (12), who found that the placement of target and flanker elements in separate hemifields produced less crowding than those in the same hemifield. Although we show significant crowding effects in our study (with clear threshold elevation and assimilation effects), we did not observe a difference based on the location of the flanker elements. We suggest that this null result arose because although the position of these elements projects to distinct hemifields, the overlap in receptive field size at the eccentricities used in both our study and that of Liu et al. would project the stimulus to both hemispheres, at least to some extent, making it difficult to separate the elements. We also found in pilot testing that the parameters used by Liu et al. gave poor visibility for several participants (see Methods), suggesting the effects may be driven more by stimulus detectability than by crowding. Our observation in Experiment 3 that radial flankers cause both greater threshold elevation and assimilation than tangential flankers offers further evidence against cortical distance as the sole predictor of crowding. Because the cortical magnification of the visual field along the radial axis leads to less compression radially than the representation along the tangential or polar angle axis (54, 58), cortical distance makes the opposite prediction to the effects we observe.

It nonetheless remains possible that cortical distance may have some influence over crowding through modulations of receptive field overlap. Predictions based on cortical distance have been shown to predict the rise in crowding with eccentricity in both peripheral (24, 31) and para-foveal (33) vision. It is of course also difficult to disentangle these effects from those of receptive field size, given that the two are negatively correlated (38). A mismatch between the two could nonetheless lead to variations in receptive field overlap – for instance, if the rise in receptive field size with eccentricity were faster than the increase in cortical distance then receptive field overlap would increase. It may be that small variations in cortical distance could underlie some of the variations in crowding, particularly if these factors were to vary in conjunction with others like receptive field size and shape to alter receptive field overlap. Our point is not that these factors are completely implausible as contributors to crowding, but rather that on their own they are insufficient to explain the entirety of the variations observed.

Receptive field overlap can also explain variations in crowding not measured in the current study. Similar to the upper-lower anisotropy, crowding is greater on the vertical meridian than the horizontal (4–6). This too could be driven by variations in receptive field overlap between the elements, in turn driven by increases in receptive field size (51) or density/spacing along these locations of the visual field. Similarly, many studies have observed an inward-outward anisotropy whereby outwards-positioned flankers (farther from fixation than the target) produce more crowding than inwards-positioned flankers (73, 74). The increase in receptive field size with eccentricity would cause the spatial distribution of responses to the outermost flankers to overlap with the target to a greater extent than the innermost flankers. Although this effect has also been attributed to cortical distance (32), others have noted differential crowding when two radially-positioned elements at fixed separation are interchangeably set as the target or flanker, despite their positions (and cortical distance) being held constant (75, 76). Receptive field overlap can explain this, since the outermost element will overlap with the inner one to a greater extent than the reverse configuration due to the size of the receptive fields involved. Finally, the increase in crowding with increases in flanker number (77) could also be driven by an increase in the overlap between the spatial distribution of the response to the flankers and that of the target. Of course, these effects do not increase indefinitely (78), making it likely that crowding does not increase beyond some asymptotic level of receptive field overlap.

Finally, the potential for receptive field overlap to drive the linked effects of crowding on appearance and performance fits with an emerging picture of crowding as an adaptive process (8, 67, 79). That is, what the visual system needs to represent an object clearly is a population response that sufficiently separates the responses of the target from background clutter. This is possible in regions near the fovea, where a large number of small receptive fields are present. In these regions, inhibitory interactions in the surround of receptive fields could even emphasise the differences between stimuli. In contrast, for peripheral representations with large receptive fields that overlap substantially, the representation of the target and flankers becomes pooled. Variations between these two extremes are then evident as the target and flanker locations vary. The push and pull between these factors thus plays a key role in determining what we can see at locations across the visual field.

## Materials and methods

### Apparatus

Experiments were programmed using MATLAB (MathWorks Ltd.) and Psychtoolbox 3 (80, 81). In Experiment 1, stimuli were presented on a LaCie Electron 22 Blue CRT monitor with 1152×870 pixel resolution and 75 Hz refresh rate. The monitor was calibrated with a Minolta LS110 photometer and linearized via look-up tables, giving a mean and maximum luminance of 50cd/m^2^ and 100cd/m^2^, respectively. Experiments 2 and 3 were presented on a Mitsubishi Diamond Plus 230SB monitor with 1400×1050 pixel resolution and a 75Hz refresh rate, and a mean and maximum luminance of 53.5 cd/m^2^ and 107 cd/m^2^, respectively. Participants were tested in a darkened room, with stimuli viewed binocularly from 50 cm and head movements minimised through a head-and-chin rest. An EyeLink 1000 desktop eye tracker (SR Research) was used to monitor fixation. Responses were made via numerical keypad.

### Participants

In Experiment 1, 10 participants were tested (8 female, aged 20-37), including two of the authors (JG and KJ). The remainder were naïve with respect to the aims of the study. In Experiment 2, 8 participants were tested (4 female, aged 20-34), including 3 of the authors (JD, JG & RF) and 5 new naïve participants. Experiment 3 also tested 8 participants (5 female, aged from 20-34), including 3 authors (JD, JG & RF) and 5 new participants. In each case, participants had normal or corrected-to-normal visual acuity. All participants were provided with an information sheet and gave written informed consent prior to starting the study. Procedures were approved by the UCL Experimental Psychology ethics committee.

### Stimuli and Procedures

#### Experiment 1

The first experiment examined whether the placement of flanker objects in the same vs. opposite hemifields would modulate both the strength of crowding (measured via threshold elevation) and its effects on appearance (measured via bias). The experiment had a 3×2×2 design, with factors for crowding condition (target alone, flanker ipsilateral, flanker contralateral), target location (left or right of the vertical meridian) and flanker orientation (clockwise or counterclockwise of the target). The latter two factors were included to counterbalance target location and orientation judgements, and were subsequently pooled to examine the effect of crowding condition.

Stimuli were Gabor elements with even phase, a spatial frequency of 2 cycles/degree, 75% Michelson contrast, and a spatial window with a standard deviation of 0.25°. These values differ from those of Liu, Jiang (12) and were selected for stimulus visibility – pilot testing revealed that the higher spatial frequency used by Liu, Jiang (12) gave poor visibility for several of our participants. In order to measure the effect of crowding on appearance, flankers were oriented Gabors with a fixed orientation offset (unlike the checkerboard Gabors used by 12). Participants fixated on a white Gaussian blob with a standard deviation of 3 arcmin, located near the bottom edge of the monitor. As in the study by Liu et al. (2009), the target could appear at one of two locations, left or right of the vertical meridian, at an eccentricity of 15° in the upper visual field. In each case, the centre-to-centre distance from the target to the vertical meridian was 1°. The target was either presented alone or in the presence of one flanker. When present, the flanker was located either left or right of the flanker, which depending on the target location corresponded to an ipsilateral or contralateral (opposite hemifield) presentation. The centre-to-centre separation between target and flanker was kept constant at 2°.

On each trial, the target was presented with one of 9 orientations around vertical (90°), ranging from 58° to 122° in steps of 8°. When present, the flanker was oriented ±10° from vertical. These values were selected during pilot testing to allow clearly measurable crowding effects for all participants. Stimuli in all conditions were presented for a duration of 200ms, after which they were replaced by a mask composed of 1/f noise, presented for a further 200ms (Figure 1B). Masks were presented within a circular envelope with a diameter of 7.9°, including a cosine edge spanning 1°, which allowed for coverage of all stimuli when centred on the target location. At the offset of the mask, participants could respond with a two-alternative forced choice (2AFC) judgement regarding whether the target was clockwise (CW) or counter-clockwise (CCW) of vertical. A 500ms inter-trial interval followed the response, with only the fixation point shown on screen. No feedback was given regarding performance.

To ensure clarity regarding which of the elements was the target and to minimise the possibility of confusion between the target and flanker elements, target and flanker location conditions were blocked separately. This was emphasised through instructions depicting which Gabor was the target element, presented at the beginning of each trial block and after any re-calibration of the EyeLink. This gave 6 separate blocks, given the 3 crowding conditions and 2 target-location conditions. Each block of trials contained 10 repeats of each target-flanker combination. This gave 90 trials for uncrowded blocks (given the 9 orientations tested) and 180 for crowded blocks (since CW and CCW flanker orientations were interleaved within blocks). These trials were preceded by five practice trials which were subsequently discarded.

Given the specificity in stimulus spacing required for these manipulations of cortical distance, the EyeLink was used to ensure steady fixation throughout stimulus presentation. Trials were cancelled when fixation strayed beyond an area of 1° radius around fixation during stimulus presentation, as well as when blinks were detected. Cancelled trials were moved to the end of the block to be repeated. As the target was kept at a constant 1° separation from the vertical meridian, this ensured that fixational shifts never allowed either target or flanker elements to move from one hemifield to the other. An average of 9.8% of trials were excluded and re-run in this manner.

Each of the 6 blocks of trials was repeated 4 times in a random order to give 3600 trials per participant (plus those cancelled by eye movements) spread over 3-4 one-hour testing periods. Before starting the test trial blocks, participants were given several practice blocks to familiarise themselves with the experiment and were not allowed to commence the test trial blocks without receiving more than 80% correct twice in a row on the practice blocks. Participants were given breaks where necessary to minimise fatigue.

Data was collected as the proportion of CCW responses and pooled across blocks, separately for each crowding condition, target location, and flanker orientation. Psychometric functions were fit to the data using a least-squared error minimisation approach, with two free parameters (slope and midpoint). To quantify these biases, we took the orientation value at which the psychometric function reached its midpoint at 50% CCW responses. To quantify performance, thresholds were taken as the difference in orientation required to shift performance from the midpoint to 75% CCW responses. Given the symmetrical effect of flanker orientation on midpoints (as in Figure 2A), we converted these bias values to assimilation scores by subtracting the 90° reference point and reversing the sign of CCW flankers.

#### Experiment 2

To examine the upper-lower anisotropy, target orientation judgements were examined either for an isolated target or with two radial flankers. Unflanked conditions had a 2×4 design, with factors for visual field (upper vs. lower), target eccentricity (from 4-16°). Flanked conditions had an additional factor for flanker orientations (clockwise or counterclockwise of the target) to give a 2×4×2 design, though the latter was then pooled across to examine the effects of visual field and eccentricity.

Stimuli were generated as in Experiment 1. Participants again fixated on a Gaussian blob, with Gabor elements presented peripherally. In the unflanked conditions, a single oriented target was presented at one of four eccentricities (4, 8, 12, or 16°) in either the upper or lower visual field. For stimuli in the lower visual field, the fixation element was shifted 200 pixels above the screen centre, with the equivalent downward shift for stimuli in the upper field. In the flanked conditions, the target was presented between two flankers positioned along the vertical meridian (i.e. the radial dimension with most crowding; 39). Flanker orientations were increased to ±22.5° from vertical, given prior demonstrations of clear variations in assimilation using these values (24). Both flankers had the same orientation on a given trial, with clockwise and counter-clockwise values interleaved in each block of trials. When unflanked, the target was presented with one of 9 orientations ranging from 78-102° in steps of 3°. When flanked this was increased to 54-126° in steps of 9°, as determined during pilot testing.

To equate visibility across eccentricity, element properties were scaled using the cortical magnification function developed by Duncan and Boynton (82), as in equation 1:

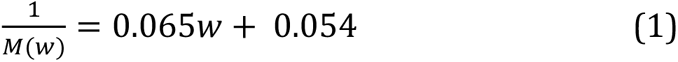

Here, *M* is the cortical magnification factor at a given eccentricity, *w*. The output of this equation was used to set the spatial frequency of the Gabor elements, giving values of 3.2, 1.7, 1.2, and 0.9 cycles/deg. across the 4 eccentricities. To maintain a constant number of cycles per element, the standard deviation of the Gaussian envelope was set as the reciprocal of the spatial frequency divided by 2. This gave 2 cycles per element in each case, with standard deviations of 0.16, 0.29, 0.42, and 0.56° for each eccentricity. The centre-to-centre separation between target and flanker elements was kept as a multiple of eccentricity, using the eccentricity multiplied by 0.3. This follows the ‘Bouma law’ whereby crowding is strongest within 0.5× the target eccentricity (1, 83), giving centre-to-centre separations of 1.2, 2.4, 3.6 and 4.8°, respectively. As before, stimuli appeared on the screen for 200ms, followed by a 1/f noise mask (scaled to cover all 3 elements at each eccentricity) for 200ms, at which point the 2AFC response was allowed.

When unflanked, each block of trials involved 10 repeats of each target orientation randomly interleaved at a single eccentricity to give 90 trials per block. With flankers, the two orientation conditions were interleaved, giving 180 trials per block. Participants repeated each block of trials 3 times, interleaved in pseudo-random order. Because the upper and lower field locations required shifts in the fixation point, the researcher adjusted the chin rest for each in order to maintain a neutral resting point for fixation (i.e. with eyes straight ahead). As a result, upper and lower conditions were clustered together, with adjustments of the chin rest in between. As in Experiment 1, participants completed practice blocks of trials prior to completing the main experiment.

#### Experiment 3

To examine the radial-tangential anisotropy, target orientation judgements were examined in the upper visual field, either for an isolated target or with two flankers on either the radial or tangential axis. Flanked conditions had a 2×5×2 design, with factors for flanker axis (radial vs. tangential), target-flanker separation (from 1.6-3.2°) and flanker orientation (clockwise or counterclockwise of the target), with the latter then pooled across to examine the effects of flanker axis and separation. Unflanked conditions were also measured to determine baseline performance.

Stimuli were again Gabors in peripheral vision whilst participants fixated on a Gaussian blob, here positioned near the bottom of the screen. The target Gabor was presented at 8 degrees eccentricity in the upper visual field, along the vertical midline. Following the calculations of Experiment 2, the spatial frequency of the Gabor was 1.7cycles/degree, with a 0.29° standard deviation of the Gaussian window.

As in Experiment 2, the unflanked target was presented with one of 9 orientations ranging from 78-102° in steps of 3°. When flanked this was increased to 54-126° in steps of 9°. Flankers were again presented with orientations of ±22.5° from vertical, with both flankers sharing the same orientation on a given trial. Target-flanker separation varied from 1.6-3.2° in steps of 0.4°, which corresponded to separations ranging from 0.2-0.4× the target eccentricity. Stimuli were presented for 200ms, followed by a 1/f mask that covered all elements for 200ms, at which point the 2AFC response was made.

When unflanked, each block of trials involved 10 repeats of each target orientation randomly interleaved to give 90 trials per block. With flankers, the two orientation conditions were interleaved, giving 180 trials per block. Each block of trials was conducted with a single target-flanker separation on either the radial or tangential axis. This gave 1 unflanked and 10 flanked configurations, each of which was repeated 3 times, interleaved randomly.

### Population models of crowding

To simulate the orientation judgements made by participants, we first simulated the responses of a population of orientation-selective neurons similar to those of cortical area V1 (84). Given the inverse relation between tuning bandwidths and the population response (85), we simulated the population response directly as a wrapped Gaussian profile of responses to orientation. This was characterised as:

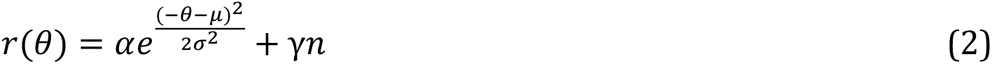

where *r*(θ) represents the population response at a given orientation θ (spanning ±90° around vertical), α the height of the population response (set to 1), μ the orientation producing the peak response, and σ the standard deviation of the Gaussian (set to give a full-width at half-maximum of 30° to approximate V1 selectivity). Gaussian noise *n* was added to this response, with a magnitude of γ (the first free parameter). Responses outside the range ±90° were wrapped by either subtracting or adding 180° to the orientation and summing the responses. Flanker population responses had the same form and the same fixed parameters, with a second free parameter for γ, representing late noise introduced by the crowding process.

To simulate errors of repulsion, inhibitory surrounds were added to the population response, as in our model of motion crowding (18), and matching both the physiology of V1 neurons (86) and physiological measures of crowding (87). Surrounds were added as a second Gaussian distribution (Equation 1), with a peak of 0.3 for the population responses to the target (given estimates that the strength of inhibition is 30-40% that of excitation; 88) and 1.0 for flankers (to be modulated by flanker weights, below), and a FWHM of 90°. This distribution was then subtracted from the excitatory Gaussian response described above.

Population responses were determined for target and flanker orientations separately and combined via weights. Variations in the effect of crowding were produced by varying these weights from 0-1, with a corresponding target weight of 1 minus the flanker weight. To simulate the effects of target eccentricity and the upper-lower anisotropy in Experiment 2, flanker weights were varied linearly over eccentricity. Separate slope parameters were fit for the lower and upper visual fields (the third and fourth free parameters), with the same intercept value used for each (the fifth and final free parameter). Given the lack of repulsion, the same weight was applied to both positive and negative components of the population response. Because the default excitatory response had a higher peak (1) than the inhibitory surround (0.3) this gave predominantly assimilative interactions within the population. The best-fitting weighting functions are shown in Figure 7A.

**Figure 7.**
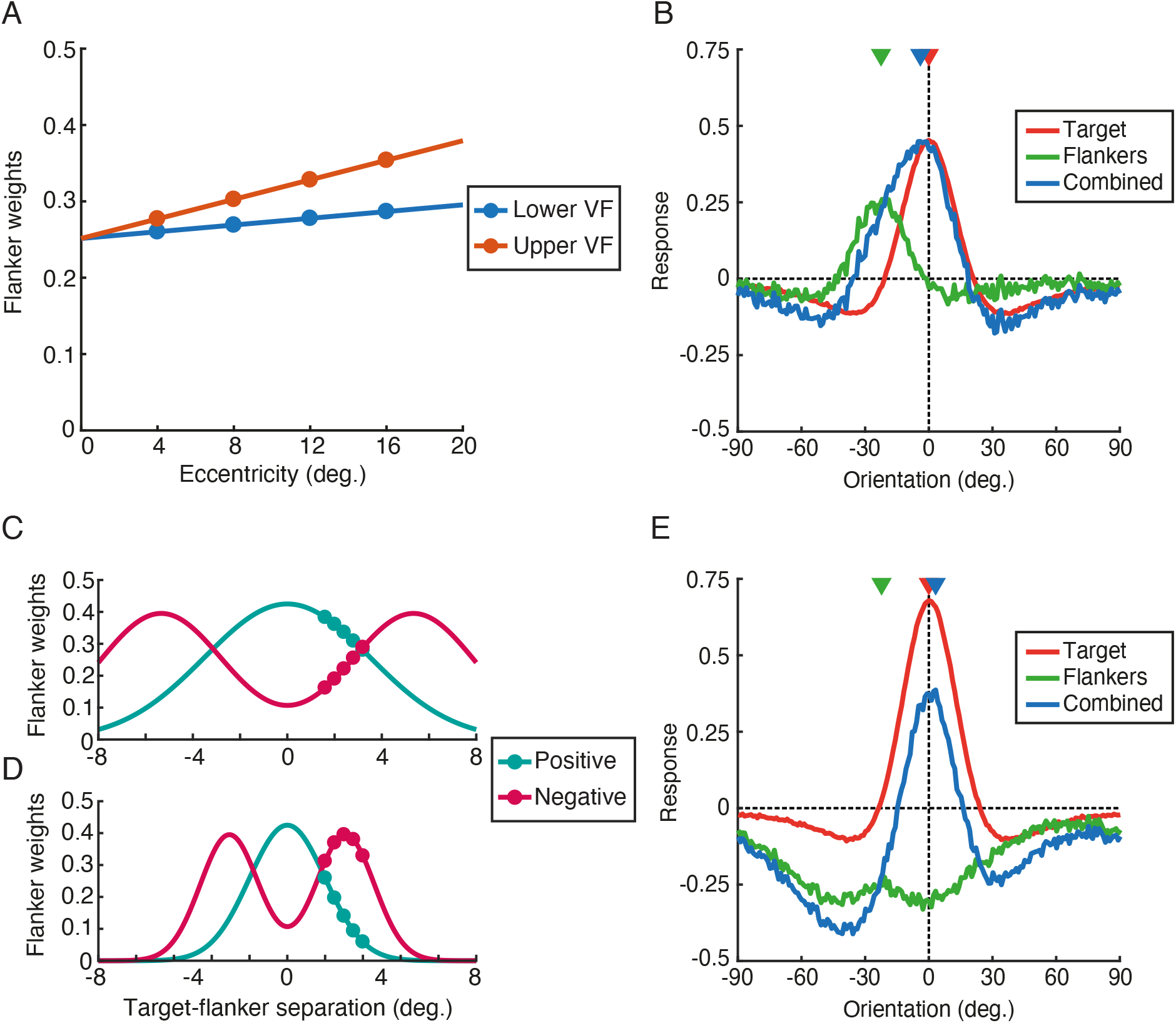
Details of the population-pooling models of Experiments 2 and 3. **A.** Flanker weights for each eccentricity tested in Experiment 2, separately for the upper (yellow) and lower (blue) visual fields. **B.** Example population responses to the target (red), flankers (blue), and combined response (yellow) for a vertically oriented target with flankers oriented 22.5° clockwise, plotted as a function of the preferred orientation of detectors on the x-axis. The veridical values of the target and flanker orientations are shown as red and blue triangles, with the peak response of the combined distribution shown as a yellow triangle. This combination gives an assimilation error. **C.** Weighting fields for orientation in Experiment 3, plotted as a function of target-flanker separation (in units of eccentricity), separately for the positive (solid) and negative (dashed) population responses. **D.** Weights on the radial dimension, plotted as in panel C. **E.** Example population responses for a vertical target with flankers oriented 22.5° clockwise at an intermediate target-flanker separation where inhibition dominates, which gives an error of repulsion. Plotting conventions as in panel B.

For a given trial of the simulated experiment, flanked responses *C* were determined as function of the orientation *θ*, with the form:

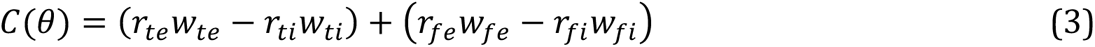

where *r_te_* was the excitatory Gaussian population response to the target (following equation 2), *r_ti_* the inhibitory response, and *r_fe_* and *r_fi_* the excitatory and inhibitory flanker responses, respectively. Weight values are denoted as *w_fe_* for the excitatory flanker values and W_fi_ as the inhibitory weight, which were selected according to the target eccentricity and visual field. For the target *w_te_* was 1-*w_fe_* and *w_ti_* was 1-*w_fi_*.

Example population distributions (averaged across 1024 trials) are shown in Figure 7B, with a vertical target and flankers oriented 22.5° counter-clockwise. Distributions of target (red line) and flanker responses (green) have had their respective weights applied. Due to the overlap in the target and flanker distributions on the clockwise side of the population, and inhibition on the counter-clockwise side, the combined sum of these responses (blue) peaks at a clockwise value intermediate between target and flanker orientations – an error of assimilation. Perceived orientation was derived in this way for each simulated trial, with the sign of the peak taken as the forced-choice response (CW/CCW of vertical). Target and flanker orientation conditions were identical to those of Experiment 2, with 1024 trials per condition in the simulation. As with the behavioural responses, percent CCW was computed for each target orientation in each flanker condition, with psychometric functions fit to determine midpoint and threshold values.

Best-fitting parameters were first determined using a coarse grid search through the parameter space to find the least squares error between the measured midpoint and threshold values and their simulated counterparts. This coarse fit was then used to seed a fine-fitting procedure using *fminsearch* in MATLAB. Best-fitting parameters were 0.251 for the intercept of the weighting function, with slopes of 0.002 and 0.006 for the lower and upper fields respectively. Early Gaussian noise was set to 0.070 and late/flanker noise to 0.486. The output of the model is plotted against the data in Figure 3 for Experiment 2.

In Experiment 3, participants showed errors of both assimilation and repulsion. Because our population used both positive and negative components, two weighting functions were used, as in our prior model (18). To capture the predominance of assimilation at close target-flanker separations and the rise in repulsion at larger separations, positive weights were determined by a unimodal Gaussian function (as in Equation 2, albeit across target-flanker separation *δ* instead of orientation *θ*). The positive weight function was plotted as a function of target-flanker separation and centred on 0, with two free parameters for the peak height and standard deviation, respectively. A bimodal Gaussian function was used for the negative weights, with the form:

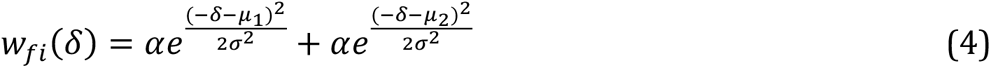

Inhibitory flanker weights (*w_fi_*) were determined as a function of target-flanker separation δ, with the peak α and standard deviation σ matched for each Gaussian and set as the third and fourth free parameters of the model. The bimodal Gaussian was centred on zero, with the peaks set at a separation (μ_2_-μ_1_) of 4 times the standard deviation of the excitatory Gaussian. This ensured the trade-off between assimilation at close separations, with a rise in repulsion at intermediate values before these weights receded at the largest target-flanker separations. The difference between radial and tangential dimensions was given by a radial-tangential multiplier (varying from 0-1; the fifth and final free parameter), applied to the standard deviation of the flanker weight function on the tangential dimension. This multiplier narrowed the range of target-flanker separations with crowding, applied equally to both positive and negative weighting functions. As before, the corresponding target weight was always 1 minus the flanker weight for both positive and negative components.

The best-fitting weighting functions are shown in Figure 7C for the radial dimension and 7D for the tangential. Example distributions (averaged across 1024 trials) with a vertical target and flankers oriented 22.5° clockwise are shown in Figure 7E. Here the flanker weights give a predominantly inhibitory population response, which causes the combined distribution to shift away from the veridical target and flanker values towards counter-clockwise orientations – a repulsion error. Best-fitting parameters, determined as above, were 0.425 for the peak of the positive weighting field, with an SD of 0.438, 0.395 for the peak of the negative weighting field, with an SD of 0.334, and a radial-tangential factor of 0.460. Early Gaussian noise was set to 0.080 and late/flanker noise to 0.448, comparable to the model for Experiment 2. The output of this model is plotted against the data for Experiment 3 in Figure 4.

### Receptive field overlap modelling

In developing our model of receptive field overlap, we first consider a simple example with a 3×3 grid of 9 neurons in the upper visual field. Each detector is a two-dimensional Gaussian element of the form:

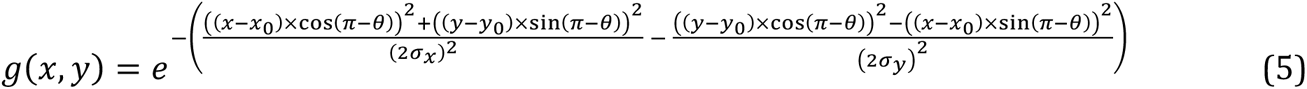

Here, the peak location of the Gaussian receptive field is given by (*x_0_*, *y_0_*), with a standard deviation on each axis set by *σ_x_* and *σ_y_*, and an orientation of the Gaussian set by *θ*. Our stimuli were Gabor elements presented within two-dimensional Gaussian contrast envelopes, meaning we can also use Equation 4 to represent the stimulus location in the visual field, with the same parameters as in our experiments. The response of each detector to a given element in our stimuli was obtained by convolving its Gaussian receptive field with the Gaussian stimulus element and taking the sum. In other words, its response was determined by the spatial overlap between the receptive field and the presented stimuli. To determine the spatial distribution of the population response to our stimuli, each detector contributed its own receptive-field profile to the final ‘image’ of the stimulus – a detector with a small receptive field would produce a tightly distributed estimate of the target location; a detector with a larger receptive field would produce a broader estimate. The magnitude of this Gaussian spatial profile was given by multiplying the Gaussian receptive field by the magnitude of the overlap between the receptive field and the stimulus (as above).

In the full model, used to simulate the variations of Experiments 2 and 3, our model population was simulated in a rectangularly spaced grid stretching to ±30° eccentricity in the upper and lower visual fields, with detectors spaced up to ±14° either side of the vertical meridian. Detectors were initially separated by 2° centre-to-centre separation. The rise in receptive field size with eccentricity was set using a linear function for the sigma parameter (in equation 5) that increased with eccentricity, with two free parameters for the intercept and slope of this function. To simulate the upper-lower anisotropy of Experiment 2, we added a factor to multiply the slope of this function in the lower visual field, causing it to rise less steeply with eccentricity than in the upper field (the third free parameter).

For the model with variations in both receptive field size and shape (aspect ratio), the degree of ellipticity was set as a fourth free parameter (in addition to the intercept and slope of the receptive-field size function and the upper-lower difference in slope), with 1 giving circular receptive fields and values greater than 1 giving ellipses with their major axis oriented along the radial dimension and a smaller extent tangentially. In the model with variations in receptive field size and density/spacing, two free parameters set the intercept and slope of this variation, as above. To produce the upper-lower anisotropy, the vertical spacing of the detectors was varied between the upper and lower visual fields, as a multiplier applied to the lower visual field as the third free parameter. A fourth free parameter similarly set the ratio between the vertical and horizontal spacing of detectors, in order to produce the radial-tangential anisotropy.

As with the population pooling model, we determined the best-fitting parameters first using a coarse grid search through the parameter space to find the least-squared error (fit to both Experiments 2 and 3 at the same time), which was used as the starting point for a fine-fitting procedure using the *fminsearch* function in MATLAB. Because the relation between the overlap sum and the flanker weights is somewhat arbitrary, we scaled the overlap sum values to range from zero to the maximum flanker weight in each experiment to place the values in a common space. For simplicity, and to allow comparison across experiments, receptive field models were fit only to the positive weights in the population pooling model (used in Experiment 3 but not Experiment 2). Best fitting parameters for all 3 models are shown in Table 1, derived from the fits to the flanker weights from the population pooling model.

**Table 1.** Best-fitting parameters for 3 population Receptive Field Overlap models. The first (RF size only) varied only parameters related to RF size: the intercept and slope of the linear RF size-by-eccentricity function, and a ratio applied to the slope in the upper vs. lower field. The second model (RF size + shape) also altered the aspect ratio of the RFs radially vs. tangentially. The third model varied RF size by eccentricity (the first two parameters) with variations in RF density/separation on the upper-lower and radial-tangential dimensions. Parameters that were fixed are shown in italics for each model.

|  | RF size only | RF size + shape | RF size + density |
| --- | --- | --- | --- |
| **Sigma intercept** | 0.803 | 1.552 | 1.421 |
| **Sigma slope** | 0.019 | 0.019 | 0.021 |
| **Upper-lower size ratio** | 0.895 | 0.783 | *1.0* |
| **Rad-tan aspect ratio** | *1.0* | 1.887 | *1.0* |
| **Upper-lower separation ratio** | *1.0* | *1.0* | 1.084 |
| **Rad-tan separation ratio** | *1.0* | *1.0* | 2.838 |

## Acknowledgements

This work was funded by a UK MRC Career Development Award (MR/K024817/1) to JG. Aspects of this work have been presented previously to the Vision Sciences Society (89) and the European Conference on Visual Perception (90).

## Appendix A: The role of collinearity in the radial-tangential anisotropy

As noted in the main text, there is a confounding variable in the explanation for the radial-tangential anisotropy observed in Experiment 3. Namely, because the orientations of our elements tended towards vertical, a change in flanker location also changed whether the flankers were collinear or parallel to the target Gabor. Prior studies have indeed found that contour alignment can modulate the strength of crowding, with particularly strong effects when flankers are close to collinear with the target and far less crowding with orthogonally oriented configurations (23, 60, 61). It may be then that our configuration with flankers oriented at ±22.5° from vertical gave greater crowding in the radial axis not simply because of the location of these flankers but rather due to their alignment with one another, in comparison to the tangential configuration where orientations were closer to being parallel.

In order to test this possibility, we examined the role of orientation on the radial-tangential anisotropies observed above for both performance and appearance. To do so, we compared the original configurations above with those that rotated the Gabor elements by 90°. In the novel conditions participants were required to judge the target orientation around the horizontal axis, either in isolation or in the presence of two flankers along either the radial or tangential axis, again with ±22.5° offsets from horizontal. This was compared with the vertical configurations tested above. If the effects of crowding were to be driven purely by contour alignment then we should see the effects of crowding on performance and appearance reversed for the horizontal configurations, given that the tangential elements are closer to being collinear than the radial elements with near-horizontal orientations. In contrast, if the effects are driven by flanker location then we should observe greater threshold elevation and assimilation for the radial elements than the tangential.

We again tested 8 participants (4 female, ages 20-34), including 3 of the authors (JD, JG, and RF). Apparatus details were identical to those of the main experiment, as were the majority of stimulus and procedural details. Elements were presented at a single target-flanker separation of 1.6°, selected to give the largest difference between the radial and tangential conditions. Participants were presented either with either a vertical (as in the main experiment) or horizontal target Gabor. When present, flankers were oriented ±22.5° relative to the target orientation. The distinct reference orientation conditions (vertical and horizontal) were run in separate blocks, all interleaved. Participants completed uncrowded blocks with 90 trials and crowded blocks with 180 trials (including CW/CCW flankers) for each orientation, each repeated three times.

As before, threshold elevation scores were calculated by dividing flanked thresholds by unflanked, with mean values shown in Figure S1A. A three-way mixed-effects ANOVA was run with axis (radial/tangential), reference orientation (vertical/horizontal) and participants as factors. This revealed a significant main effect of axis, *F*(1,7) = 14.92, *p* = .006, whereby radial flankers produced greater threshold elevation than tangential in both target orientation conditions. Although there is a trend towards higher threshold elevation with the horizontal reference orientation, the main effect of reference orientation was not significant, *F*(1,7) = 3.31, *p* = 0.112, nor was the main effect of participant, *F*(7,7) = 2.55, *p* = .120. No interactions were significant (all *F*<2).

**Figure S1.**
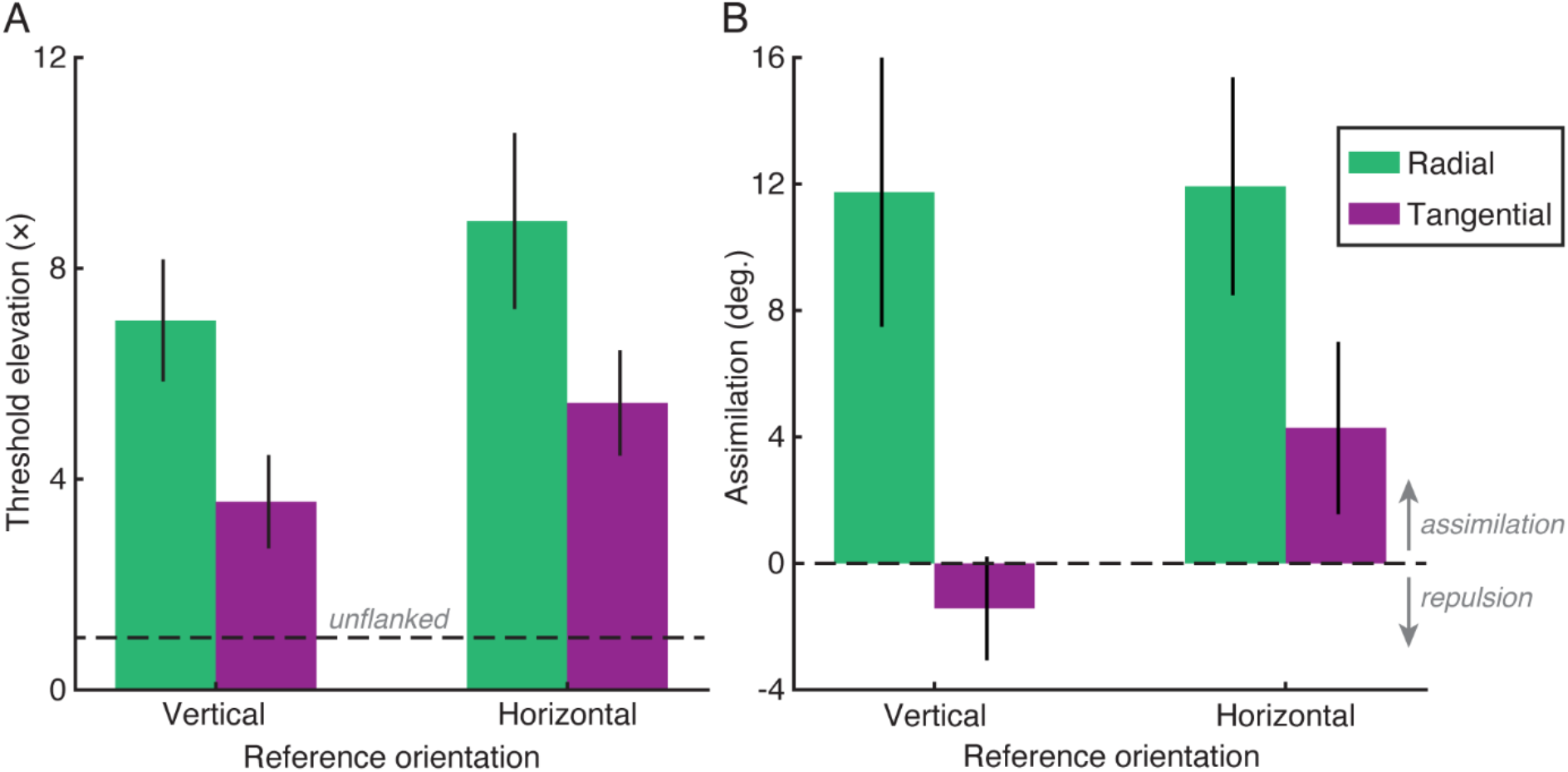
Results from the control experiment on the role of collinearity in Experiment 3. **A.** Mean threshold elevation scores for flankers on the radial (green) and tangential (purple) axes, separately for Gabors oriented around a vertical reference (left) or horizontal reference (right). Unflanked performance is indicated via the dashed black line. Error bars show the SEM. Lines show the output of the best-fitting population pooling model. **B.** Mean assimilation scores, where positive values indicate assimilation and negative values repulsion (separated by the dashed line), plotted as in panel A.

Assimilation values varied similarly, as shown in Figure S1B. Again the main effect of axis was significant, *F*(1,7) = 15.61, *p* = .006, with radial flankers producing greater assimilation than tangential in both target orientation conditions. Although there are some differences evident for the two reference orientations, e.g. with repulsion evident for radial but not tangential flankers, the main effect of target orientation was not significant, *F*(1,7) = 2.11, *p* = .190, nor was the main effect of participant, *F*(7,7) = 4.82, *p* = .247. No interactions were significant (all *F*<2).

Although there are slight differences between the two reference orientation conditions, the clearest effect on both performance and appearance is driven by the placement of the flankers on the radial or tangential axis with respect to fixation. Radial flankers produce higher threshold elevation and greater assimilation than tangential flankers, which produce some repulsion and lower threshold elevation. The small differences observed may relate to the radial bias in human vision (91), where sensitivity to orientation is greatest for elements oriented orthogonally to the radial axis, or to collinearity effects driven by contour integration processes (23, 60, 61). However, this is clearly not the driving factor behind the effects observed.

## Appendix B: The correlation between assimilation and threshold elevation

A key motivation for this study was to examine whether crowded effects on performance and appearance are linked. Across the experiments reported herein, we see similar patterns of variation between assimilation and threshold elevation. Here we also examine their relatedness by examining whether the two are correlated. Figure S2A plots the results from Experiment 1 (examining hemifield effects), with threshold elevation plotted against assimilation for each participant in each flanked condition. The two are highly correlated, such that higher threshold elevation is associated with greater assimilation: r(18) = 0.608, p=0.004.

**Figure S2.**
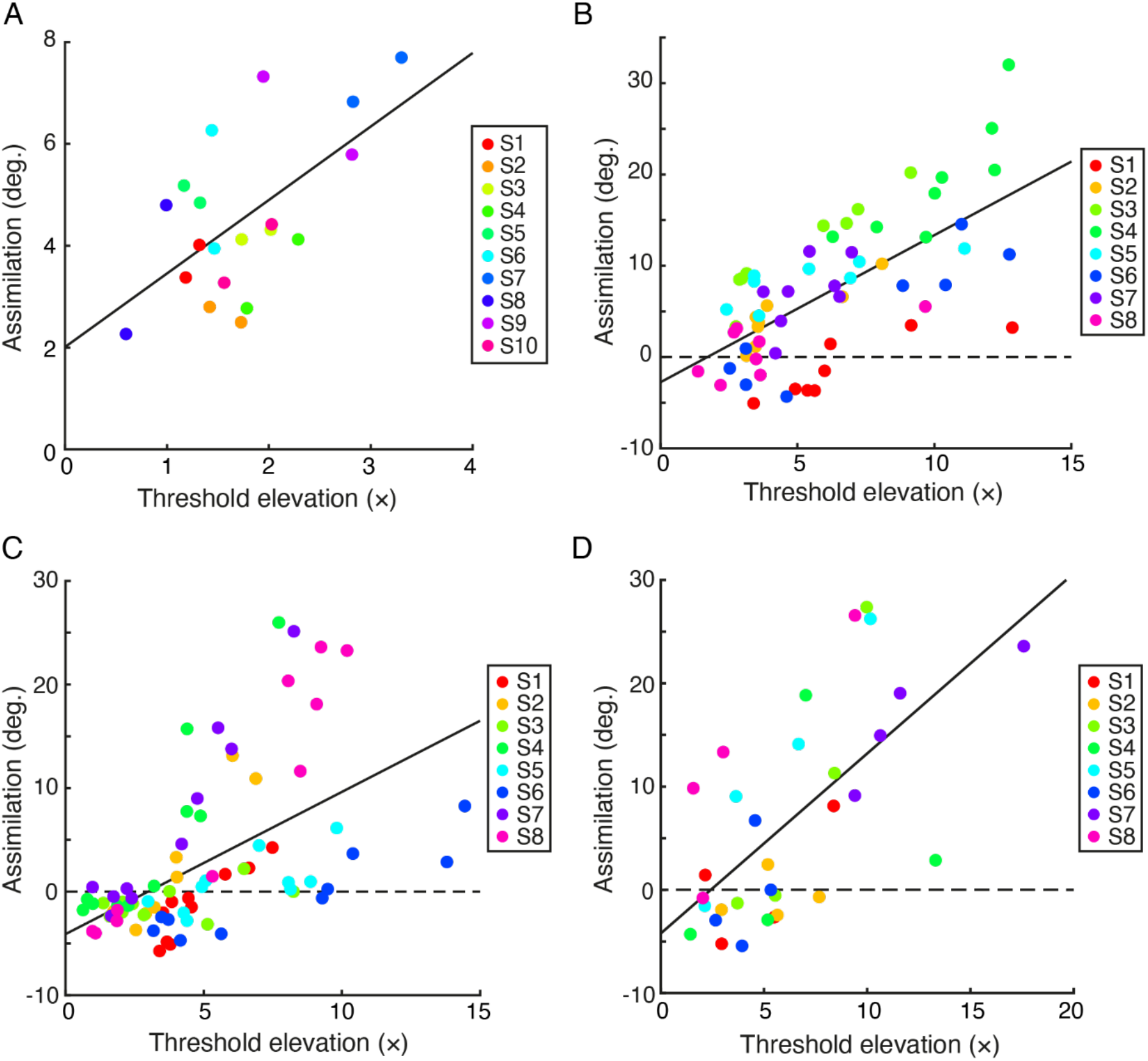
Correlations between threshold elevation and assimilation. **A.** Values from the flanked conditions of Experiment 1, with each colour showing a separate participant (see legend). **B.** Values from Experiment 2, plotted as in panel A. **C.** Values from Experiment 3. **D.** Values from Experiment S1.

This relationship is also seen in Experiment 2 (examining the upper-lower anisotropy), as plotted in Figure S2B, with a correlation of r(62) = 0.668, p<0.001. A similar correlation is observed in Experiment 3 (measuring the radial-tangential anisotropy), with r(78) = 0.554, p<0.001. Finally, in Experiment S1 (measuring collinearity effects in the radial-tangential anisotropy), the correlation is r(30) = 0.653, p<0.001. These correlations between crowded effects on performance and appearance are consistent with arguments for a general link between bias and discriminability (63), though the linear correlations that we observe suggest a more direct link between the two, as others have suggested (64).

